# Co-segregation of recombinant chromatids maintains genome-wide heterozygosity in an asexual nematode

**DOI:** 10.1101/2023.03.17.533182

**Authors:** Caroline Blanc, Nathanaelle Saclier, Ehouarn Le Faou, Lucas Marie-Orleach, Eva Wenger, Celian Diblasi, Sylvain Glemin, Nicolas Galtier, Marie Delattre

## Abstract

In asexual animals, female meiosis is modified to produce diploid oocytes. Associated with recombination, this is expected to lead to a rapid loss of heterozygosity, with adverse effects on fitness. Many asexuals, however, have a heterozygous genome, the underlying mechanisms being most often unknown. Cytological and population genomic analyses in the nematode *Mesorhabditis belari* revealed another case of recombining asexual being highly heterozygous genome-wide. We demonstrated that heterozygosity is maintained because the recombinant chromatids of each chromosome pair co-segregate during the unique meiotic division. A theoretical model confirmed that this segregation bias is necessary to account for the observed pattern and likely to evolve under a wide range of conditions. Our study uncovers a new type of cell division involving Directed Chromatid Assortment.

**One sentence summary:** Genome wide heterozygosity in the asexual nematode *Mesorhabditis belari* is achieved by directed assortment of recombinant chromatids during female meiosis

## Introduction

Asexual animal species are composed of females, which produce diploid daughters without paternal genome contribution. Asexuality requires the production of diploid oocytes and hence, a modified female meiosis. Asexuality, which is derived from sexuality, has emerged multiple times and independently over the course of evolution and many routes to producing diploid oocytes have been documented (*1–3*).

Depending on the type of cellular modification, different genetic outcomes are expected (Figure 1A). A common prediction is that most modifications should lead to loss of heterozygosity (LOH). For some species, the entire meiotic program is achieved as in sexual species, however the final haploid nucleus undergoes a duplication event (gamete duplication), which immediately generates a homozygous individual (Figure 1A). When there is homologous recombination and one of the two meiotic divisions fails, either because the division is abortive or because the product of meiosis fuse back, LOH is also expected either distally or proximally to the crossover location (Figure 1A). Hence, maintenance of heterozygosity is theoretically expected only in species for which meiotic recombination is largely reduced or totally abolished (*4*).

**Figure 1:**
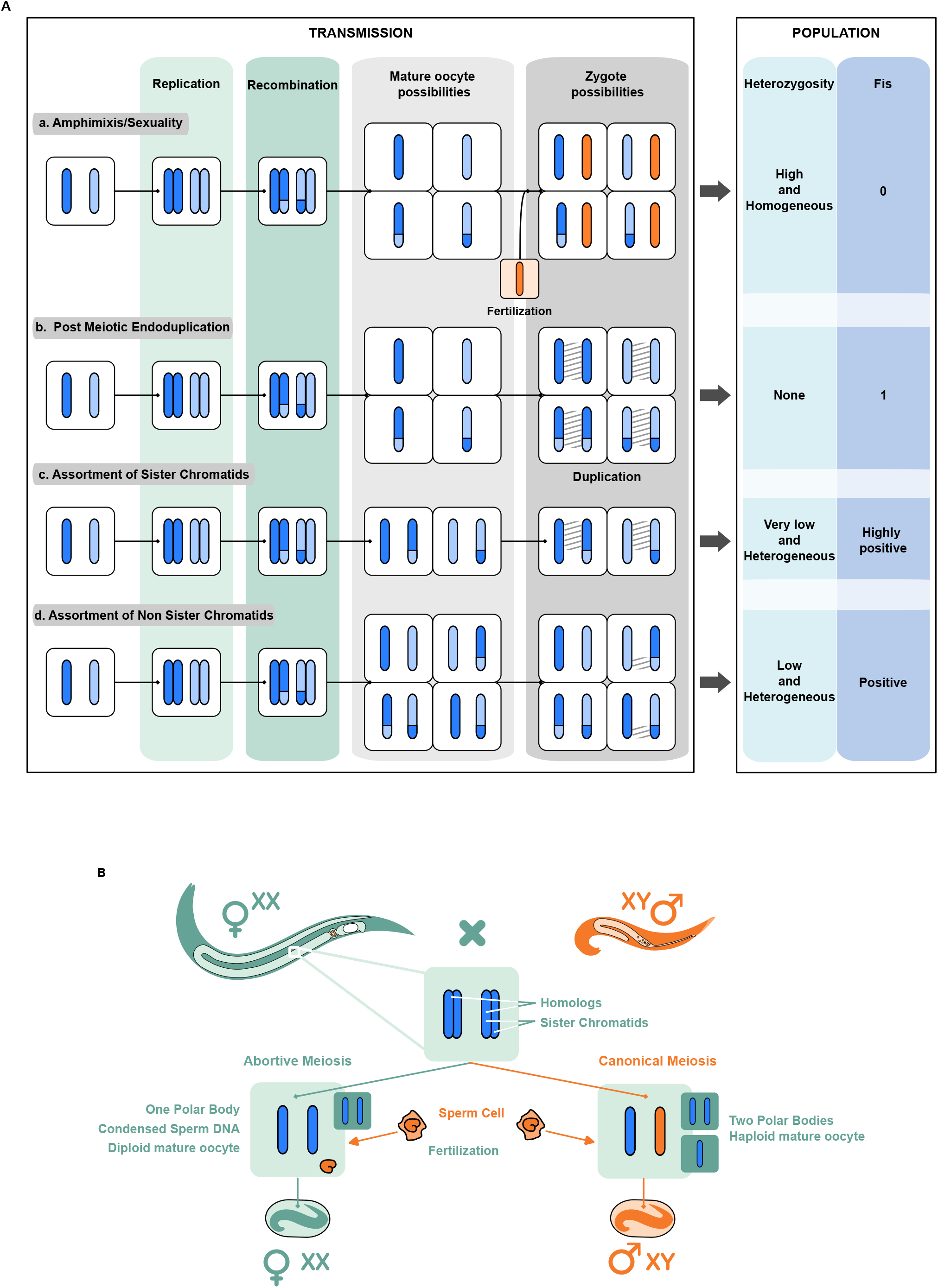
Genetic expectation upon modification of meiosis in asexuals and reproductive system of *Mesorhabditis belari*. A) Description of the expected genetic composition of a zygote, and of a population, upon different types of modifications of the meiotic program (b-d). Canonical meiotic division as found in regular sexual species is shown on the top (a). Each bar represents a chromatid. The paternal chromatid is in orange. The maternal chromatids are in dark and light blue, corresponding to homologous chromosomes. Upon recombination, assortment of chromatids after modification of meiosis generate stretches of homozygosity, shown with hatching. Consequently, the level of heterozygosity in the population (right column) is decreased. The coefficient of inbreeding (Fis) corresponds to the expected level of heterozygosity compared to Hardy-Weinberg expectation (in a randomly mating population). B) Schematic representation of the reproductive system found in *M. belari* as described in (*13*). Females (in green) produce two types of oocytes. Through canonical meiosis, a single chromatid per chromosome is transmitted to the oocyte (in blue). The sperm provides in single chromatid (in orange). The resulting diploid individuals give rise to males. 90% of the oocytes are however diploid (incomplete meiosis, on the left) in which case the sperm does not contribute DNA. The individuals give rise to females.

LOH is expected to negatively affect fitness due to the exposure of recessive deleterious mutations at diploid state. This has been suggested as a potential cause of the relative scarcity of asexual organisms in nature (*5*), and a selective pressure for reduced recombination rates in asexuals (*6*). Species that maintain some level of heterozygosity do not expose deleterious mutations and as such may circumvent some of the drawbacks of asexuality. In agreement with this expectation, a number of asexual lineages display appreciable amounts of heterozygosity (*7–10*). Yet, except in species for which a total loss of recombination has been demonstrated (for instance (*11, 12*)), the mechanisms of heterozygosity maintenance are still debated (*7*). Therefore, there is still a need to confront the cytological description and empirical genome data to reach a clear understanding of the genomic and cellular constraints in asexual animals.

We explored the mechanism of meiosis in the auto-pseudogamous nematode *Mesorhabditis belari* (Figure 1B) (*13*). In this species, a female produces a majority of diploid oocytes, which although fertilized by a sperm, develop only from the maternal DNA and become females (gynogenetic embryos). The same female also produces ~10% of haploid oocytes through regular meiosis, which once fertilized undergo fusion of the parental genomes. These amphimictic diploid embryos will give rise to males because active sperm cells always carry a Y chromosome (*13*). Hence, this species produces 90% asexual females and 10% sexual males. In *M. belari* females, which most likely have maintained recombination for the production of regular oocytes (for the rare males), we asked which modification of meiosis has been selected to produce the asexual females and with which genomic consequence for the species.

## Results

### Diploid oocytes of M. belari are formed after failure of the first meiotic division

We first asked which steps of meiosis were modified to produce diploid oocytes in *M. belari*. We followed meiosis, making use of the spatio-temporal organization of the gonad, as found in the well-studied *Caenorhabditis elegans* species and other Rhabditidae nematodes (Figure 2A). We had previously shown that this species is diploid and carries 2n=20 holocentric chromosomes (*13*). First, analysis of oocytes in diakinesis revealed that the 20 chromosomes were always paired into 10 units (n > 200 oocytes). Moreover, many bivalents had a crossed-shape structure (Figure 2A), which resemble the chiasmata of holocentric chromosomes found in *C. elegans (14, 15*), strongly suggesting that chromosomes undergo homologous recombination in *M. belari*.

**Figure 2:**
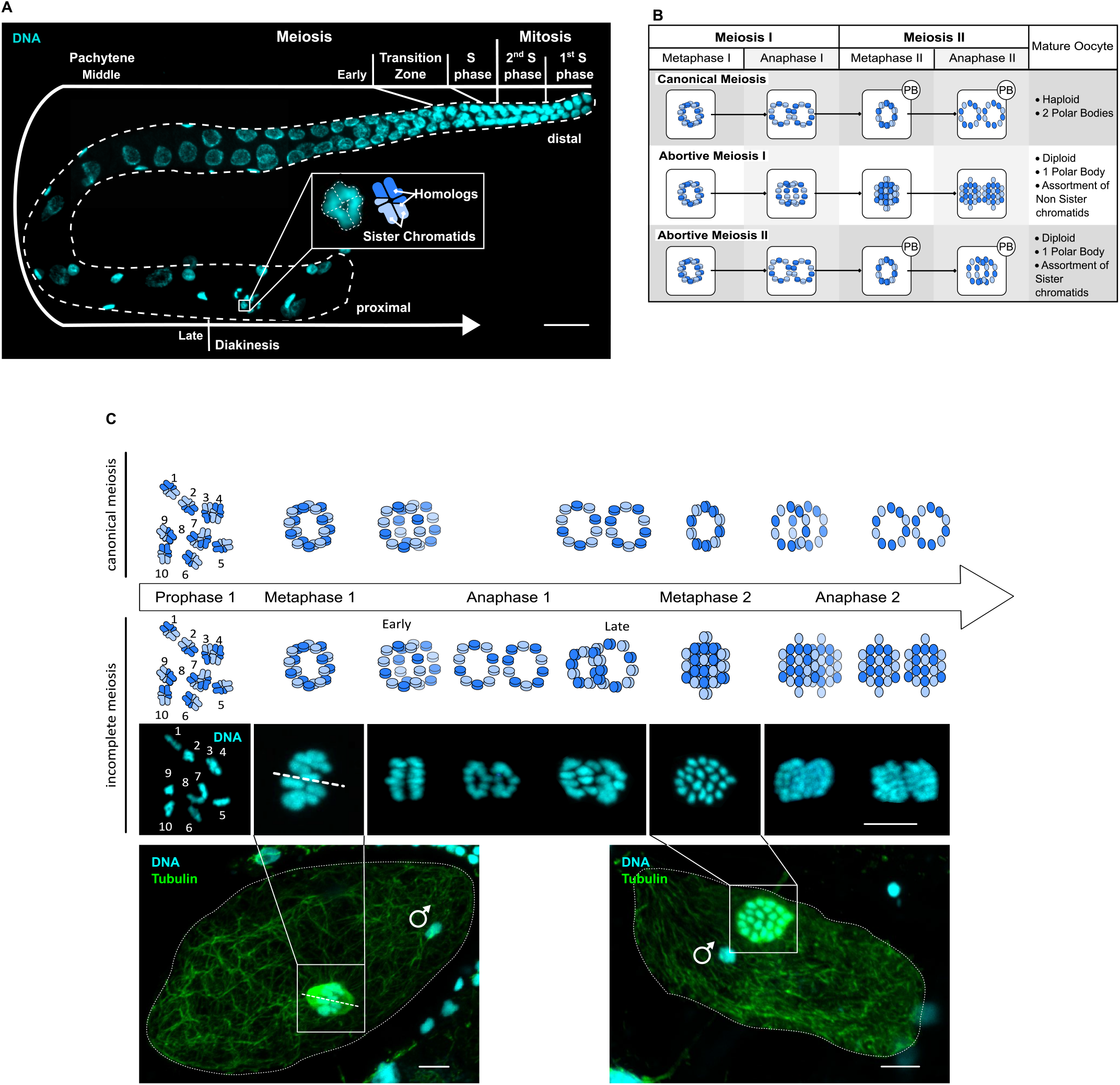
Cytological evidence of abortive meiosis I in *M. belari* females. A) Gonad of a *M. belari* female stained for DNA, showing the progression of meiotic cells along the tract. Oocytes in diakinesis are found on the distal part, with chiasmatic chromosomes. DNA is in blue. Scale bar is 20 μm. The holocentric bivalent chromosomes show a typical cross shape. The bivalents are schematized in blue, one chromosome is in dark blue and its homolog is shown in light blue. B) Expected figures of chromosome organization upon failure of meiosis I or meiosis II in *M. belari*. A canonical meiosis is shown on the top. At metaphase, chromosomes orient as a ring. During anaphase, chromosomes (Anaphase I) or chromatids (Anaphase II) segregate as two rings. PB represent the polar body. C) Reconstitution of *M. belari* meiosis in amphimictic (canonical meiosis) and gynogenetic (incomplete meiosis) embryos from fixed samples. On the bottom, representative gynogenetic embryos, from which the images are taken, are show. A sperm DNA is visible although it will remain condensed and will fuse with the female DNA. Tubulin is in green and DNA is in blue. *M. belari* is diploid carrying 2=20 chromosomes. The dotted line represents the long axis of the meiotic spindle along which the bivalent chromosomes align. Scale bar is 5 μm.

Next, we reconstructed the different steps of meiotic divisions (Figure 2B, C). We found, as is the case for *C. elegans*, that oocytes are arrested at prometaphase of meiosis I and meiosis resumes after fertilization. The long axis of the bivalents is oriented parallel to the spindle axis, as expected for holocentric chromosomes in regular meiosis (Figure 2C) (*14*). In the Rhabditidae sexual species studied so far, a polar body is extruded at each meiotic division. The polar bodies are easily recognizable as tiny cells at the edge of the embryo (*16*). We had previously shown that all gynogenetic embryos in *M. belari* had only one polar body demonstrating that one meiotic division is suppressed (*13*). We reasoned that if meiosis I was abortive, the 10 bivalents should dissociate into 20 univalents and no polar body should be detected at this stage. These univalents should next enter anaphase of meiosis II, showing two times 20 DNA stained bodies (Figure 2B). On the contrary, if meiosis I was successful, a polar body would be extruded, and the 10 univalents would disassemble into 20 units corresponding to 20 sister chromatids after failure of meiosis II (Figure 2B). We found many cells showing a metaphase plates containing 20 DNA stained bodies. We also detected anaphase figures with two times 20 DNA stained bodies (Figure 2C). Importantly, none of these cells had produced a polar body. Of note, we also found images of metaphase with only 10 DNA stained bodies, which we interpret as being either figures of regular meiosis (~10% are expected) or the initial step of meiosis I before abortion. From these results, we concluded that diploid oocytes in *M. belari* are formed after failure of the first anaphase of meiosis I. This modification will lead to the assortment of non-sister chromatids in the oocytes (equational division only). In the presence of recombination for all chromosomes, this pattern of inheritance should progressively lead to LOH, distally to the crossing-over.

### M. belari has a widely heterozygous genome

We analyzed the level and the distribution of heterozygosity in the genome of *M. belari* females, from our lab strain JU2817 and nine other wild strains, which had been sampled in different locations around the world (*17*). We sequenced mixed stage animals from each strain and mapped the short reads on the assembled genome of *M. belari* JU2817. Genome-wide heterozygosity was computed by counting the proportion of heterozygous positions relative to the total number of positions using ANGSD (*18*). Each strain being isofemale (see Material and Methods), the genotype of a strain corresponds to the genotype of a single individual.

We found that all ten strains had approximately the same level of heterozygosity of about 1,3% [sd = 0.2] (*i.e*. one residue every 75nt is heterozygous), demonstrating that the strains behaved similarly in the wild and in the lab (Figure S1 and Table S1). This level of heterozygosity is unexpectedly high for a meiotic asexual experiencing regular recombination; it is 10 times as high as in the self-fertilizing nematode *Caenorhabditis elegans* (*19*) and similar to natural populations of the fruitfly *Drosophila melanogaster* (*20*), for instance. Heterozygosity could be maintained in most parts of the chromosomes and lost only in subtelomeric regions if crossing-overs were restricted to chromosome ends but we found that heterozygosity was uniform along all contigs (Figure S1).

To further confirm the maintenance of heterozygosity, we performed a genome-wide analysis of genotype inheritance, from mother to daughters in *M. belari* JU2817. We performed this analysis on the transcriptome which can be easily obtained from single worms. We analyzed three female individuals descended from the same mother. RNAseq reads were mapped to the previously assembled *M. belari* JU2817 transcriptome, genotypes were called, focusing on sufficiently covered contigs and positions. Under the assumption of active recombination and random segregation of chromatids, large chromosomal segments - and therefore a substantial number of SNPs and contigs - are expected to be homozygous in some of the females (Table 1). In contrast to this prediction, we found only a small minority of SNPs for which at least one daughter had a homozygous genotype (Table 1). These SNPs were most often surrounded by SNPs, for which all three daughters were heterozygous, suggesting there was no real stretch of homozygosity in any female. Among the >3200 contigs with more than two SNPs, only 9 contigs carried > 2 SNPs homozygous in the same daughter. Similar proportions were found when the same analysis was performed in another autospeudogamous species, *M. monhystera*. As a control, we also analyzed two sexual *Mesorhabditis* species *M. longespiculosa* and *M. spiculigera*. In these species, we found a large fraction of SNPs with homozygous genotypes (Table 1), as expected under random mating of gametes, with 737 (out of 1423, ~52%) and 1371 (out of 2953, ~46%) contigs, respectively, carrying >2 SNPs homozygous in the same daughter. This analysis indicates that the modified meiosis in autopseudogamous species of *Mesorhabditis* (almost) entirely preserves heterozygosity, from one generation to the next, and in the population overtime.

**Table 1.**
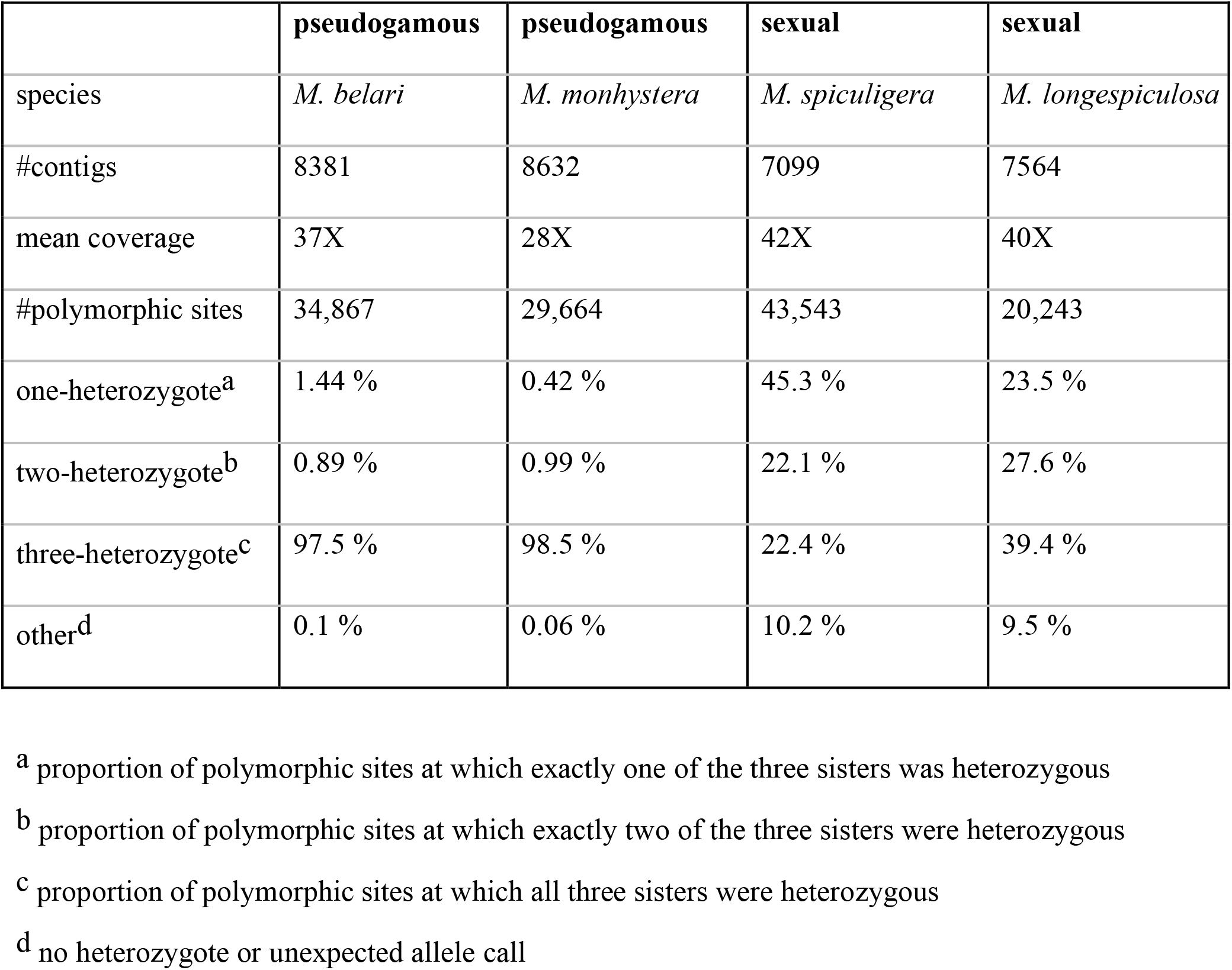
Patterns of shared heterozygosity among sisters in four species of *Mesorhabditis*.

These results seem to contradict our initial cytological observations. We therefore asked if despite the presence of structures, which resemble chiasmata, homologous recombination might be absent, which would then explain the maintenance of heterozygosity in the short and long term upon assortment of non-sister chromatid.

### All homologous chromosomes recombine during female meiosis

We wished to directly visualize crossing-over as a formal proof that homologous recombination occurs during *M. belari* female meiosis. To this end, we used the thymidine analog EdU, which is incorporated into replicating DNA during oogenesis and can be fluorescently labelled. *M. belari* females were bathed in EdU (pulse phase) and then allowed to recover (chase phase) so that cells next divided and replicated without EdU. We optimized the pulse and chase periods to obtain chromosomes harboring only one EdU-labelled chromatid (Figure S2). Upon recombination, we then expected a strain exchange between one EdU-labelled (shown in pink in Figure 3) and one non EdU-labelled chromatid (shown in blue in Figure 3, labelled with Hoechst) in 50% of cases, generating bicolor chromatids (blue/pink) (Figure S2, Figure 3). We first analyzed the color of chromatids in diakinesis oocytes, in which homologous chromosomes form chiasmata, *i.e*. bivalents. The expected figures of crossing-over in holocentric chromosomes has been described in (*21*) and is depicted in Figure 3A. From 8 oocytes, we identified 26 bivalent chromosomes whose orientation allowed us to unambiguously distinguish the chromatids within the chiasma. For 12 of them, the two opposed chromatids had the same color and could not be analyzed. Among the 14 showing opposed chromatids of different colors, 13 bivalents, showed an exchange of chromatids and only one showed no exchange (Figure 3A). We also analyzed chromosomes in the female pronuclei of gynogenetic eggs during the first or second cell-cycle, when chromosomes are condensed and chromatids clearly visible. Cycles of DNA replication and mitosis had occurred in the absence of EdU in embryos, generating many chromatids devoid of EdU. Nevertheless, we counted 49 bicolor chromatids from 12 embryos (one representative embryo shown in Figure 3B). This analysis also revealed that chromatid exchange is not restricted to chromosome ends, as many chromosomes show large portions of EdU positive chromatids (Figure 3B). These results demonstrate that exchanges of strands between homologous chromosomes during meiosis are frequent, and that recombination- not restricted to telomeric regions- does occur in *M. belari* during the production of unreduced oocytes.

**Figure 3:**
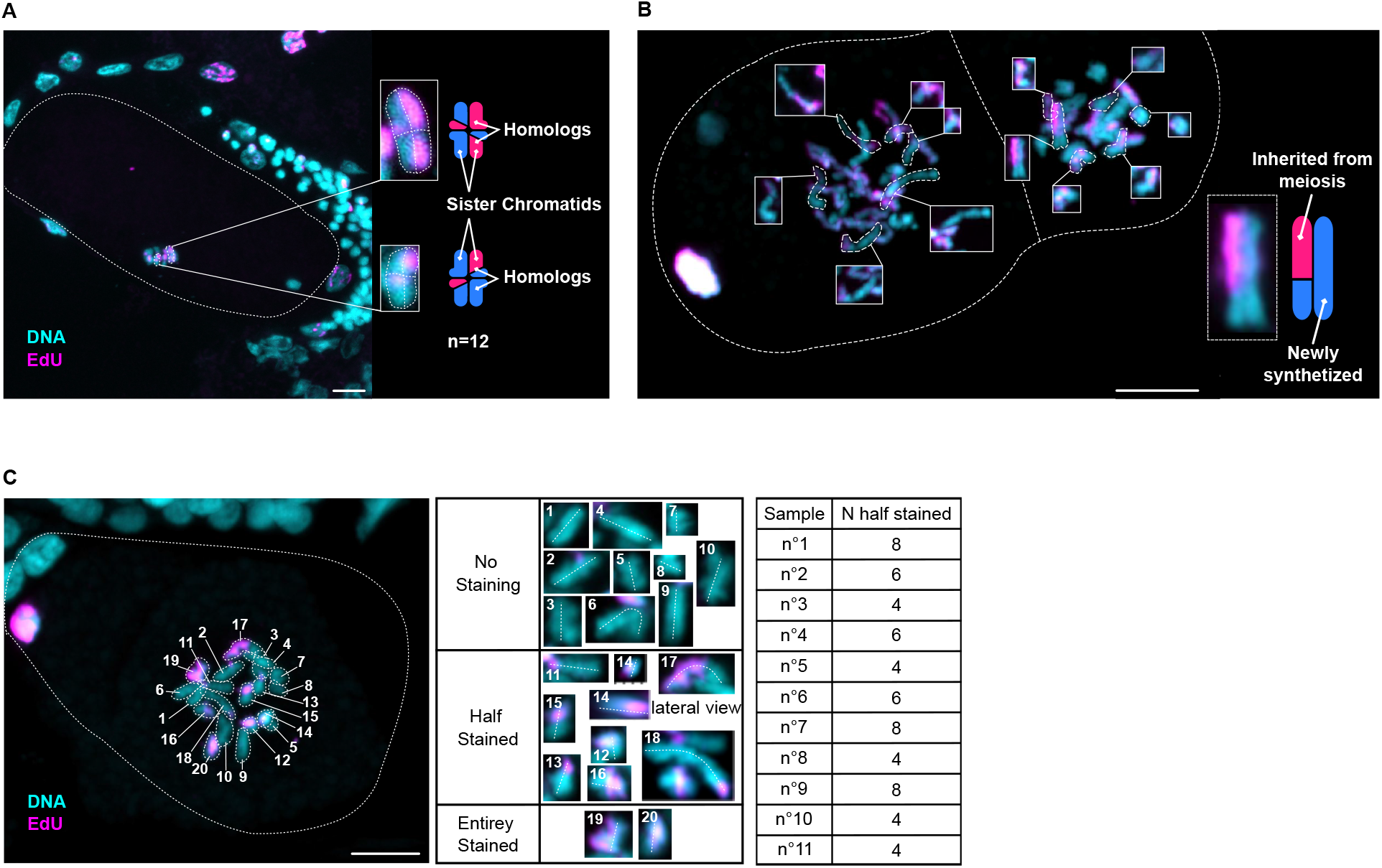
Evidence of recombination and Directed Chromatid Assortment. Fixed embryos after an EdU experiment, where one chromatid out of the two is stained with EdU (in pink). DNA is in blue. Scale bar is 5 μm. A) Embryo in metaphase of meiosis I (*i.e*. no polar body is extruded). The metaphase plate is perpendicular to the glass slide. Few chromosomes are visible on this lateral view. The expected exchange of chromatids, as described in (*21*) is shown as well as the actual images in the inset. B) Two-cell stage embryo during prometaphase. Most chromosomes have a bicolor chromatid (half blue/half pink, demonstrating recombination) and an unlabeled chromatid because it has replicated during the previous S phase in the absence of EdU (entirely blue). C) One representative one-cell embryo in prometaphase. All 20 chromosomes are shown in the insets. 8 chromosomes are bicolor (#14 is shown twice). The table summarizes the count of recombinant chromatids from eleven embryos (also shown in Figure S3).

As another evidence that recombination is maintained in *M. belari*, we analyzed patterns of linkage disequilibrium (LD) across loci. In the absence of sex and recombination, alleles are expected to be strongly associated among loci, with haplotype blocks extending over long stretches of DNA. Recombination, if at work, breaks allele associations, leading to a decay of LD as the physical distance between SNPs increases (*22*). To assess the extent of LD in *M. belari*, we analyzed the genomic data previously obtained from the ten strains. We called SNPs and used the LDHelmet program to estimate the genome-wide distribution of the effective population recombination rate ρ. This analysis indicated that in both species the estimated ρ was homogeneously high across the genome (Figure S1), with a point estimate of the average ρ of 0.038 per base pair. This implies that LD is lost as the distance between the considered loci exceeds ~100 bp. The average estimated ρ in *M. belari* was similar to estimates reported in sexual species of arthropods, such as *Drosophila melanogaster* (*20*), and indicative of a high effective population recombination rate in this auto-pseudogamous species.

### Chromatid segregation is biased during the unique meiotic division of M. belari embryos

Our cytological and genomic data are contradictory because recombination and random assortment of non-sister chromatids should lead to LOH. We reasoned that maintenance of heterozygosity from mother to daughters, and at the population level, can be achieved despite recombination, if either the two recombinants, or the two non-recombinant chromatids of a given chromosome pair co-segregate into the egg during the unique division of meiosis. We validated this hypothesis using our EdU experiment. Because *M. belari* chromosomes cannot be distinguished cytologically (*i.e*. pairs cannot be recognized), we used a statistical approach. We reasoned that under the hypothesis of co-segregating recombinant chromatids, we should always find an even number of recombinant chromatids in the nuclei of one-cell stage embryos, before the first mitosis. Such a pattern would be obtained very rarely in the case of random segregation of the 20 chromatids (p-value 0.00048, binomial test, n=11, p=0.5). We reanalyzed a new set of one-cell embryos, selecting only those in which chromosomes were well spread out at prometaphase, so that chromosome axis was unambiguously identified. In the 11 embryos analyzed, we always found an even number of bicolor chromatids, ranging from 4 to 8 (Figure 3C and Figure S3). This result strongly supported that the single meiotic division of *M. belari* females is unique, as it leads to Directed Chromatid Assortment (DCA).

### Modeling the reproductive strategy of M. belari and Directed Chromatid Assortment during meiosis

To assess whether DCA could reconcile cytological and genomic data, we developed a population genetics model. As output parameters, we considered the level of heterozygosity and the Linkage Disequilibrium LD (as measured above) but also the inbreeding coefficient Fis. In a randomly mating species (Hardy-Weinberg expectation), Fis equal 0. Fis is negative if there is an excess of heterozygosity compared to the Hardy-Weinberg expectation, as found in asexuals that have lost recombination, and positive if there is a deficit of heterozygosity, as in selfers for instance (*23, 24*). By analyzing the heterozygosity found in the 10 independent wild strains of *M. belari*, we revealed a Fis close to 0, *i.e*. Fis=0.019 throughout the genome. Such a value was unexpected for a species with a rate of sexual reproduction close to 0. We had previously shown that out of 1000 females, no sexual females were produced (*13*).

We modeled the life cycle of *M. belari*, including the production of sexual males and asexual females with a biased sex ratio, the inbreeding mating structure (brother-sister mating or strong family structure as proposed by (*13*)), and the modified meiosis with variable segregation bias (Figure 4A-B). We also allowed for rare production of sexual females. Although no sexual females have been observed under laboratory conditions, they may exist at low rate in natural populations and it is an intermediate stage that necessarily occurred in the transition from sexuality to asexuality. We looked for conditions that could explain the observed genomic pattern: high heterozygosity, Fis~0, low LD.

**Figure 4:**
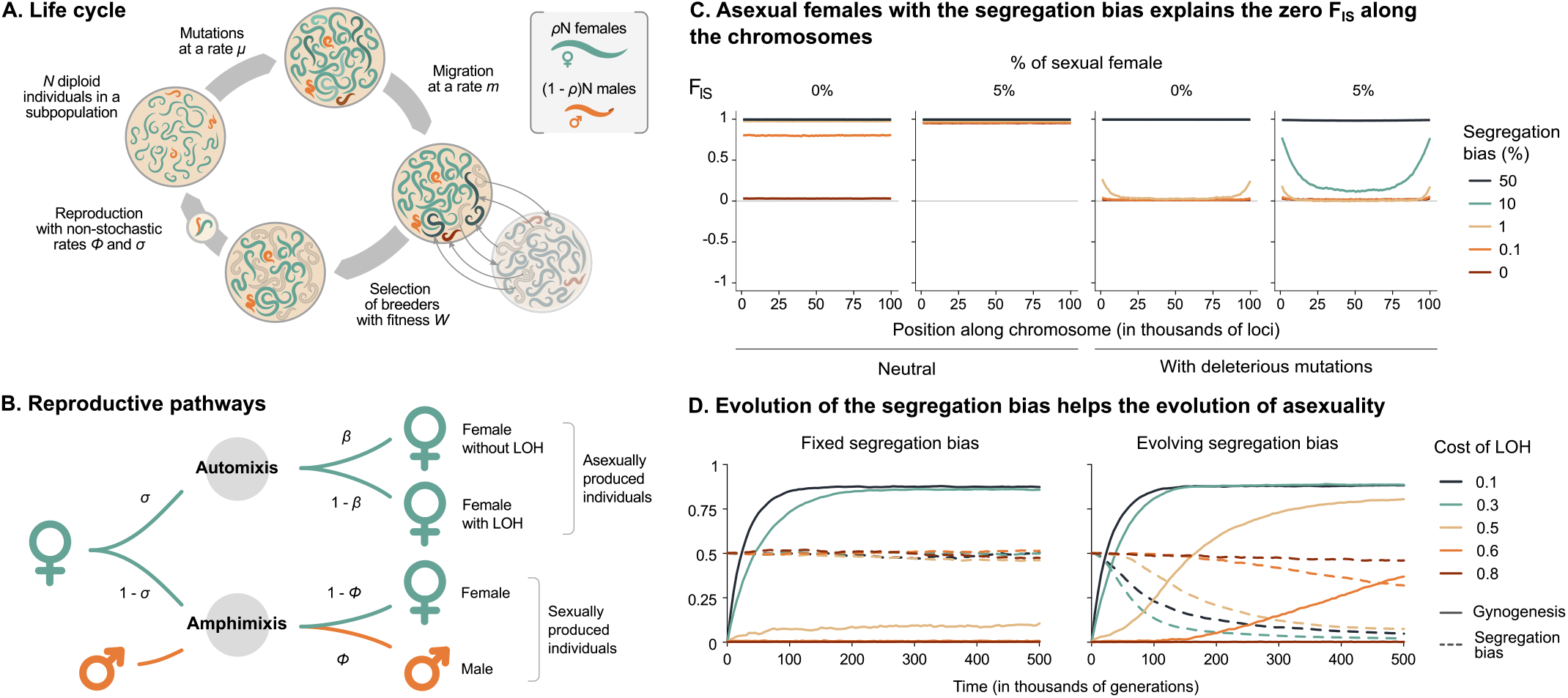
Modeling the genomic consequences and evolution of DCA in *M. belari*. A) Simulated life cycle with associated parameters. The generations are discrete and non-overlapping and we assumed an island model to simulate the high level of consanguineous mating of natural population of *M. belari*: the lower the local population size (N) and the migration rate (m), the higher the level of consanguineous mating. B) Reproduction can occur either by automixis at rate s or sexually by amphimixis at rate 1 – s. During automixis, the rate of LOH is b. b = ½ corresponds to random assortment of chromatids whereas b = 0 corresponds to strict DCA. Under sexual reproduction, females are produced at rate f f is close to zero in natural population. s and f determines the sex-ratio, r, in the population. C) Fis along a chromosome as a function of the level of consanguineous mating, with or without sexually produced females and with or without deleterious mutations. The total population size is 1000 and the migration rate is *m* = 0.0005. *N* = 1000 corresponds to a single population, *N* = 200 to five large breeding groups, and *N* = 20 to 50 small breeding groups. D) Evolution of pseudogamy from an initially sexual population with different costs of LOH when the segregation bias is fixed or allowed to evolve.

First, analytical results and multilocus simulations confirmed that DCA is needed to explain the absence of LOH. Second, considering deleterious mutations throughout the genome broaden the conditions that can explain the observed genomic patterns: strict DCA is not required and sexual females can be produced at low rate (Figure 4C and Supp. Text). Actually, the production of sexual females at very low rate better explains the low and flat LD pattern than pure asexuality (Supp. Text). The comparison of results without and with deleterious mutations also illustrates the central role that recessive deleterious mutations likely play in the system. Highly homozygotes individuals that should be produced by imperfect DCA or leaky sex (that should lead to Fis > 0, Figure 4C neutral) are selected against, maintaining Fis close to zero for a large range of conditions (Figure 4C with deleterious mutations). This also supports the idea that LOH should be costly and suggests that DCA could be selected as a LOH-preventing mechanism.

We tested this hypothesis via a modification of the initial model whereby the proportion of biased chromatid segregation can evolve. We first simulated a sexual species with no segregation bias, and added a locus controlling the proportion of asexual females produced. We assumed one mandatory crossover per chromosome. Hence, the asexual females experienced LOH with associated fitness reduction due to the expression of recessive deleterious mutations in homozygotes, a form of inbreeding depression. If inbreeding depression was higher than 0.5 it compensated for the advantage of not producing males and pseudogamy could not evolve. If inbreeding depression was lower than 0.5, pseudogamy rapidly evolved. Next, we introduced mutations at a second locus controlling DCA during asexual meiosis (in both directions: recombinants could be more positively or more negatively associated than at random). We found that mutations leading to positive association between recombinants were selected for and that the population rapidly evolved towards complete DCA, preventing the deleterious effects of LOH. Interestingly, we also found that when mutations affecting pseudogamy and chromatid assortment were introduced at the same time, the two mechanisms co-evolved, speeding up and broadening the conditions for the evolution of pseudogamy (Figure 4 and Supp. Text). On the one hand, the occurrence of some asexual females enabled the evolution of DCA. On the other hand, once DCA started to evolve, it partly prevented the deleterious effect of LOH, favoring the evolution of pseudogamy, even when inbreeding depression was higher than 0.5 (Figure 4D).

Our modeling approach thus confirmed that all a priori contradictory observations can be reconciled by the mechanism of DCA during the unique meiotic division of females and proposed a selective explanation for the evolution of such a peculiar mechanism from a sexual ancestor.

## Discussion

In this study, we found that *M. belari* asexual females are produced in the presence of recombination and assortment of non-sister chromatids, which should lead to rapid LOH, distally to the crossing-over. Our genomic analysis, however, revealed a surprisingly high level of heterozygosity throughout the genome and no sign of LOH, even locally. Using a combination of cytological, genomic and modeling approaches, we demonstrated that this pattern is possible provided the recombinant chromatids of each chromosome pair are not randomly assorted but instead co-segregate during the unique meiotic division. We named this new type of non-Mendelian inheritance Directed Chromatid Assortment (DCA). With DCA, specific pairs of chromatids are chosen during cell division such that the whole set of maternal alleles is transmitted to offspring (Figure 5).

**Figure 5:**
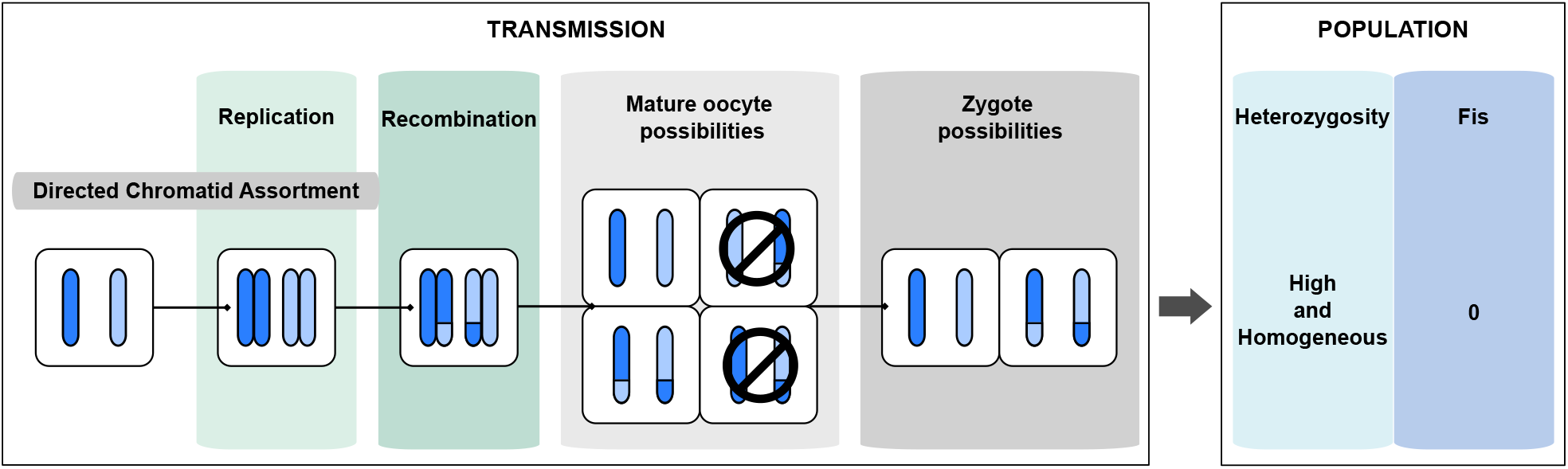
Mechanism of heterozygosity maintenance via DCA in *M. belari*. Schematic representation of meiosis and DCA during the production of diploid oocytes in *M. belari*. Homologous chromosomes (in blue) are initially heterozygous in the mother (shown in nuances of blue). After recombination and failure of meiosis I, non-sister chromatids do not segregate randomly during meiosis II. Instead, co-segregation of recombinant chromatids (in orange) maintains heterozygosity in the progeny and in the population.

Most asexual animal models have been characterized either by cytology or genomics, rarely both, whereas such a combination of approaches was here decisive. We suggest that DCA may exist in other asexuals displaying highly heterozygous genomes. For instance, DCA is compatible with the absence or very reduced LOH found in the recombining parthenogenetic water flea *Daphnia magna* (*25*), the asexual females produced by Cape Honey Bee workers (*26*) or the Rotifer *Adineta vaga* (*9*).

During their reproductive life time, *M. belari* females produce 10% reduced oocytes via regular meiotic divisions, which develop into sexual males, males being essential for sperm-dependent parthenogenesis (*13*). Hence, the meiotic program is intact in this species. This is a constraint for the evolution of asexuality because upon recombination, asexual females should experience LOH. We discovered DCA, as a new mechanism of LOH avoidance despite recombination. Another mechanism of LOH avoidance have been previously proposed for recombining asexuals, which relies on distal crossing-over location (*6, 26*). However, distal crossing-overs are very unstable for the *C. elegans* holocentric chromosomes and often lead to aneuploidy (*27*), suggesting that such mechanism of LOH avoidance could not have been selected in a holocentric species such as *M. belari*. A mechanism involving inverted meiosis (which is compatible with holocentricity), failed meiosis II and biased chromatid segregation has been proposed for the maintenance of heterozygosity in Oribatid mites (*28, 29*). Although such biased segregation of chromatids remains hypothetical in the absence of further cytological description, it is conceptually similar to the DCA we describe in our study.

How did DCA emerge mechanistically during the evolution of auto-pseudogamy in *Mesorhabditis*? One hypothesis is that the abortion of meiosis I was first fixed in the population. If sexual females were initially still produced, it compensated for LOH in the asexuals. Once DCA appeared, LOH was prevented, next allowing the loss of most sexual females. Alternatively, a modification of the meiotic program could have generated simultaneously a defect in anaphase I and a biased segregation of chromatids. At this stage, it is difficult to speculate on a molecular mechanism, and on a sequence of events, because to our knowledge no such phenotype has been described in mutants of model species. Yet, it is not inconceivable that some of the proteins that are loaded on chromatids during recombination remain specifically attached to the recombinant chromatids, contributing to a directed orientation during division, or that chromosomes experience an incomplete resolution of the crossing-overs. More work on the mechanistic basis of meiosis in *M. belari* is required to address this question.

## Funding

This work has been supported by a grant from the ANR-19-CE02-0012-01 to MD, NG and SG and PhD fellowship from CNRS to CB.

## Authors contributions

MD, NG and SG designed the experiments. SG designed the model. ELF developed the simulation code, run and analyzed the simulations with LMO and SG. CD, NS and NG analyzed the genomic data. CB, EW and MD performed the experimental work. NG, SG and MD wrote the manuscript. We acknowledge the contribution of the imaging plateform PLATIM from SFR Biosciences Lyon, the sequencing plateform Genomeast at IGBMC Strasbourg. We also thank Carine Rey for help with the treatment of the NGS raw data. We thank Raphaelle Dubruille and Benjamin Loppin for critical reading of the manuscript.

## Competing interests

All the authors declare having no competing interests.

## Data and materials availability

all data is available in the manuscript or the supplementary materials except data related to genome sequencing, which can be found on ebi.ac.uk/ena accession number PRJEB30104)

## Supplementary Materials

### Materials and Methods

#### Nematode strains and culture

*Mesorhabditis* species are maintained at 20°C on NGM plates seeded with *E. coli* OP50, following *C. elegans* protocols, as described in (*13*).

#### Immunostainings on gonads and embryos

Immunostainings were performed as described in (*13*). Gravid females were dissected on slides coated with 0.25%poly-lysine in 0,5X M9. After freeze-cracking, fixation of samples was performed by immersing slides into methanol at −20°C during at least 5 min. We used a mouse anti-tubulin antibody as primary antibody (Sigma DM1A, 1:2000) and an Alexa488 donkey anti-mouse secondary antibody (Jackson ImmunoResearch, #715-545-150, 1:2000). Both antibodies were incubated at room temperature for 45 min. DNA was stained using Hoechst 33258 at 0,5 ul/ml (Merck Sigma-Aldrich, #94403). Images were acquired using a confocal microscope (Oil immersion 63X objective – LSM800 and LSM980 Airyscan, Zeiss). Z-stacks of embryos were acquired every 0.15 μm. Finally, acquired images were treated using the ImageJ 1.53t software.

#### EdU Pulse/Chase protocol

The protocol was adapted from (*15*). In order to obtain diakinesis oocytes and early embryos for which only one chromatid per chromosome was labeled with EdU, we had to optimize the protocol for *M. belari*. First, we synchronized worms using axenization, as described in (*13*). Briefly, worms were collected and treated with bleach and NaOH in order to dissolve all individuals except the embryos. The embryo pellet was then washed and placed on plates without bacteria, allowing L1 larvae to hatch. Without food, all L1s were arrested at the same stage after 2 days. L1s were then placed back on food and allowed to grow for 72h at 20°C. At this stage, young L4 synchronized worms were collected and washed in 1X PBS, 0.1% Triton X. A pellet of ~300ul of worms was transferred into 200 ul of 10 mM EdU diluted in water (ThermoFisher A10044) to obtain a final concentration of 4 mlM EdU. The tube was transferred on a tube rotator for 4h at room temperature. After washes in M9, animals were plated onto fresh NGM plates seeded with *E. Coli* and placed at 25°C for 48 before embryos were collected for fixation. As summarized in Figure S2, we deducted that two rounds of S phase precede meiotic prophase during *M. belari* oogenesis.

#### EdU Click-it labeling

Cytology was performed following the instructions provided by the EdU Click-it kit (ThermoFisher C10337). Embryos were collected after axenization, as described above. Embryos were placed on poly-lysine coated slides, freeze-cracked and fixed in −20°C methanol. Samples were incubated with BSA 2% for 20 min at room temperature. Slides were then washed twice with 1X PBS. Following the instructions of the kit, slides were washed for 30 min at room temperature in 1X PBS with 1% Triton X (v/v) and labeled with Alexa 488-azide for 30 min at room temperature. Samples were then washed twice with 1X PBS and were incubated in a tank with Hoechst 33258 for 20 min at room temperature. Finally, slides were mounted using ProLong™ Diamond Antifade Mountant (ThermoFisher P36965) and sealed with nail polish.

#### Airyscan and ImageJ 3D analysis of EdU-labelled recombinant chromatids

As with immunostaining, confocal airyscan images were acquired using the Zeiss LSM800 Airyscan and LSM980 Airyscan using a 63x oil objective and 0.15μm interval between slides. Images were subsequently processed by the airyscan processing method (3D analysis, automatic low stringency, Zen Blue 3.3). Processed images were treated using ImageJ 1.53t software. IAnalyses of each chromosome were done using a combination of Z projection and 3D projection on the Y-axis using the brightest point method and interpolation.

#### DNA and RNA preparation for sequencing

We performed DNAseq on 10 strains of the auto-pseudogamous species *M. belari* coming from different locations in Europe. For each strain, one gravid female was initially collected in the wild and left to lay eggs in a Petri dish. This constituted a single strain, which was frozen in our collection. For sequencing, we amplified the animals and extracted the DNA for each strain. Briefly, mixed stage worms were collected, washed in M9 and a pellet of ~300 ul of worms was frozen in liquid nitrogen. After thawing, 600 ul of Cell Lysis Buffer (Qiagen Cell Lysis Solution #158906) was added, as well as 6 ul of proteinase K at 17 ug/ul and incubated for 3h at 65°C. We next incubated the mix for 1h at 37°C, supplemented with 40 ul of RNAseA (at 5 mg/ml). 200 ul of Protein Precipitation Solution (Qiagen #158912) was next added and after 5min on ice, the mix was centrifuged for 10 min at 13000 rpm at 4°C. 600 ul of isopropanol was added to the supernatant. After 10 min at room temperature, the mix was centrifuged at maximum speed and the pellet was rinsed twice in ethanol 70°C. The pellet was dried and resuspended in nuclease free water.

For the analysis of genotype inheritance in sisters, we isolated gravid females, for each species, let they lay eggs and after few days isolated 3 virgin daughters. The mRNAs of each single female were extracted using the SmartSeq2 protocol, as described in (*30*).

For all samples, genomic libraries (insert sizes of ~550 bp) were prepared using TruSeqNano and the libraries were sequenced on a HiSeq4000 with 100 bp paired-end read length.

#### Heterozygosity analysis

For each of the 10 strains of *M. belari*, reads were mapped to the assembled genome of *M. belari* with BWA (*31*). BAM files were produced with SAMtools (*32*) and heterozygosity was estimated for each strain using ANGSD (*18*) using the SFS estimation for a single sample. Recombination at the extremities of the chromosomes could explain a limited decrease in heterozygosity, the rest of the chromosome remaining non-recombining. To test this, heterozygosity was calculated on 5000 bp windows. A decrease in heterozygosity at the ends of the contigs was then looked for, graphically. At first, we detected homozygous portions in the JU2817 strain. But these portions were twice as low in coverage as the rest of the contigs, which could be explained by an assembly error due to too much divergence between the two alleles. A similarity search with blastn allowed to detect the presence of indels between the two alleles, preventing a unique assembly of these regions. This pattern was not found in the other strains and therefore explains the lower heterozygosity of this strain compared to the others (*i.e*. haplotype divergence).

#### Genotype inheritance

RNAseq was performed for three sisters in each of the two auto-pseudogamous species *M. belari* and *M. monhystera* and the two sexual species *M. spiculigera* and *M. longespiculosa*. For each species, reads from the three individuals where pooled to assemble a transcriptome: adapters were clipped from the sequences, low-quality read ends were trimmed (phred score <30) and low quality reads were discarded (remaining length <36bp) using trimmomatic (v0.39 (*33*)). Paired-end transcriptomes were de novo assembled using Trinity v2.13.2 (*34*).

Reads were mapped on their respective assembled transcriptome with BWA (*31*), BAM files were produced with SAMtools (*32*) and SNPs were called using reads2SNP (*35*) focusing on sufficiently covered contigs (minimum contig average coverage = 15X) and positions (minimum = 20X). In each species, we selected positions in which not only the called genotypes but also read frequencies varied significantly among sisters. Specifically, for each position, two multinomial models were fitted to read counts. Model M0 (three degrees of freedom) assumed a common frequency of A, C, G and T in the three sisters. Model M1 (9 degrees of freedom) rather allowed each of the three sisters to have its own frequencies of A, C, G and T reads. A likelihood ratio test was performed and we only positions in which M0 was rejected (p-val<1.e-8). This was intended to exclude positions for which the genotype varied among sisters due to uncertainty in genotype calling.

#### Measure of linkage disequilibrium

To test for the existence of recombination, we estimated the linkage disequilibrium using the ten strains of *M. belari*. For each of the 10 strains reads were mapped to the assembled genome with BWA (*31*). BAM files were produced with SAMtools (*32*) and SNPs were called with reads2SNP (*35*). To phase haplotypes, we first used WhatsHap (*36*) to extract the phase information contained in reads. Phasing was then completed using Beagle V5.3 (*37*). Linkage disequilibrium was computed on 5000 pb windows with LDhelmet V1.10 (*20*) using recommended parameters.

**Table S1:** Measure of heterozygosity in 10 strains of *M. belari*.

| Strain | Country | Location | Heterozygosity |
| --- | --- | --- | --- |
| JU2856 | France | Angles-sur-l'Anglin, Indre | 0,01381808 |
| JU2859 | United Kingdom | Cambridge | 0,01531046 |
| JU2817 | France | Orsay | 0,00735625 |
| JU3129 | United Kingdom | Coventry | 0,01419700 |
| JU3151 | France | Saint-Aigny, Indre | 0,01280673 |
| JU3152 | France | Saint-Aigny, Indre | 0,01240229 |
| JU3157 | Germany | Heidelberg | 0,01400762 |
| JU3158 | Germany | Heidelberg | 0,01366134 |
| JU3159 | Germany | Postdam | 0,01459582 |
| JU3388 | Ireland | Inis Mor, Aran Islands | 0,01361719 |

**Figure Supplementary 1:**
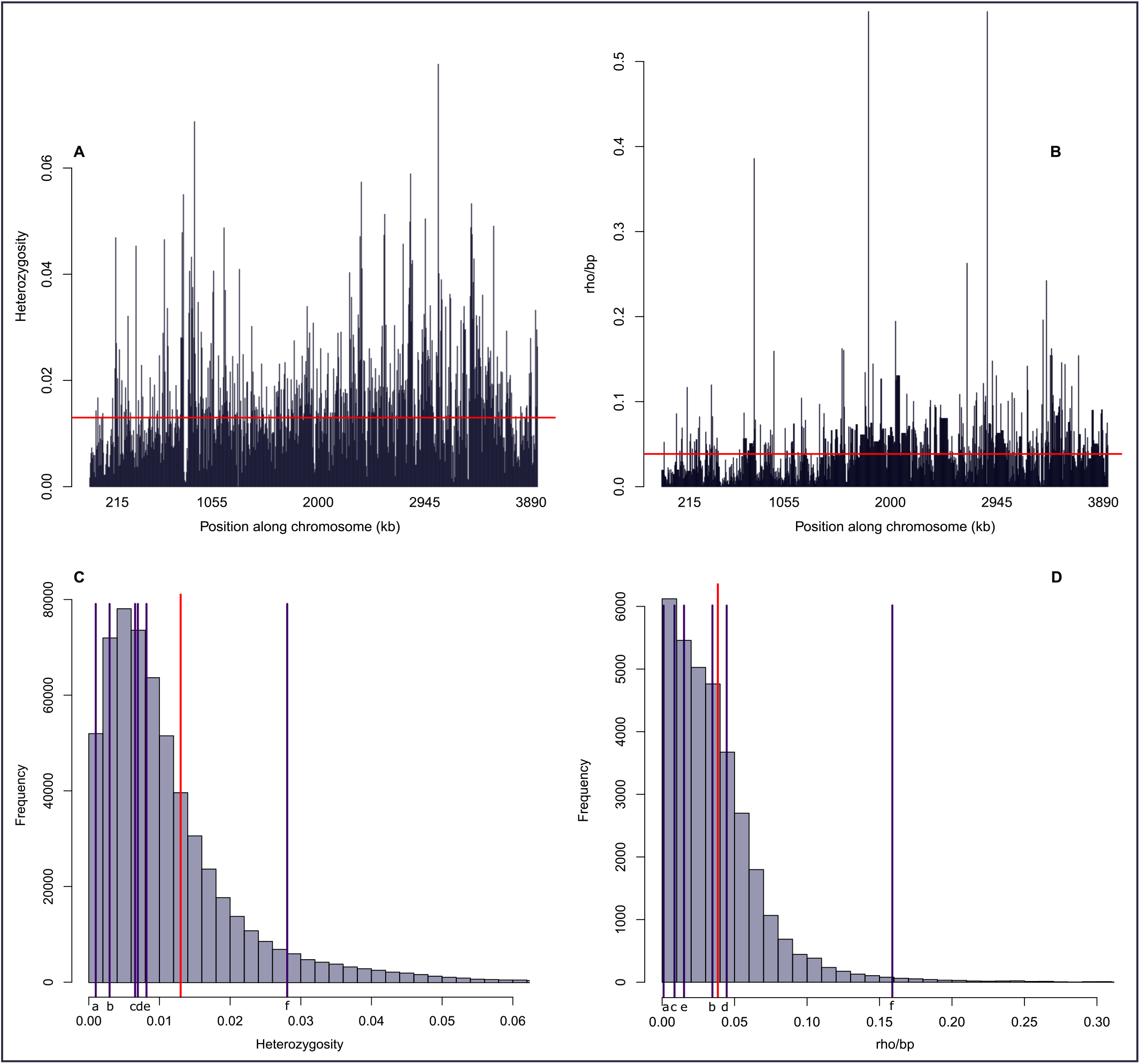
Quantification of heterozygosity and estimation of ρ. Heterozygosity (A) and recombination (B) estimated along a representative contig (contig 110) and distribution of heterozygosity (C) and recombination rates (D) computed on 5000bp windows. Genome-wide average heterozygosity and recombination rate ρ are represented by a red bar. Other sexual species, taken from the literature, are depicted with a blue bar: a) *Homo sapiens* (*38, 39*); b) *Gasterosteus aculeatus* (*40*); c) *Mus musculus* (*41, 42*); d) *Drosophila melanogaster* (*20*); e) *Taenopygia guttata* (*43*); f) *Heliconius melpomene* (*44, 45*).

**Figure Supplementary 2:**
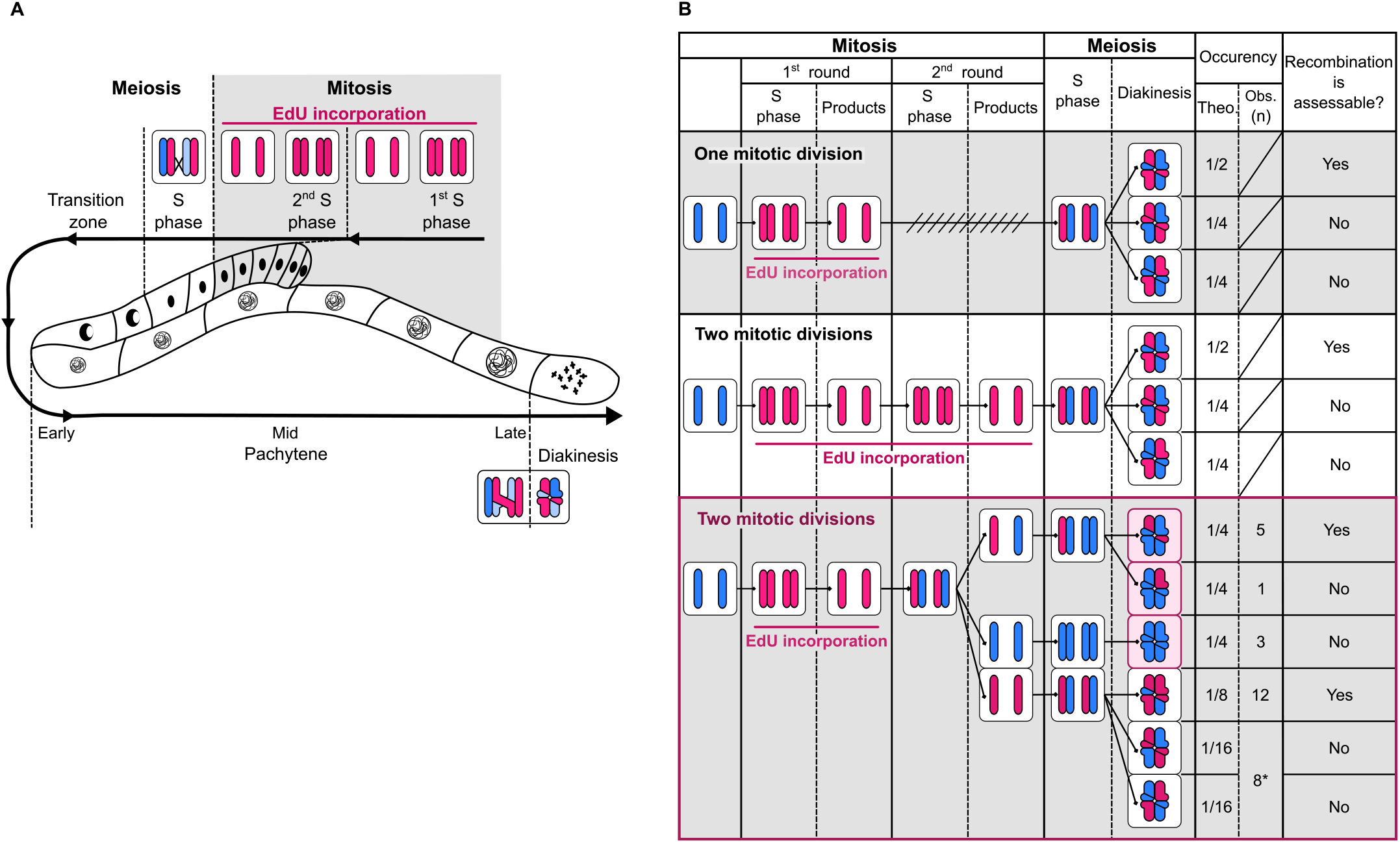
Design and expectations for the Edu experiment. A) Design of EdU pulse-chase method. Sketch of *M. belari* gonad showing the number of homologs present at each stage of oogenesis and optimal EdU incorporation during mitotic S phase (replication phase). B) Expectations for EdU labelling on the bivalents according to the number of miotic S phases and the duration of EdU incorporation (pulse phase). An account of the different EdU labels is provided on the right. EdU is pink and DNA is blue. The observed pattern of EdU incorporation is consistent with EdU exposure during 2 rounds of S phase.

**Figure Supplementary 3:**
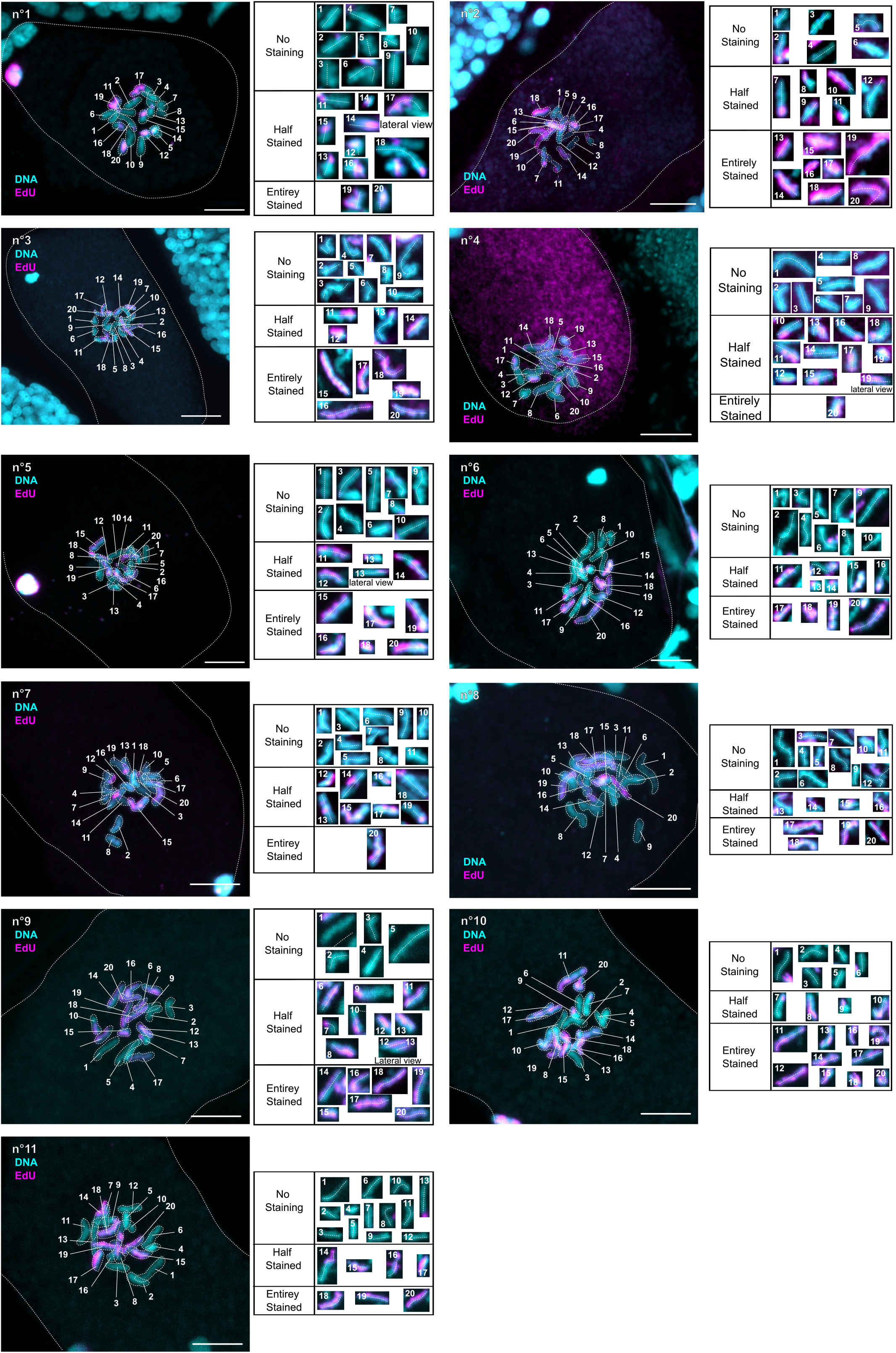
Evidence of Directed Chromatid Assortment. 11 images of fixed embryos at one-cell stage. EdU is in pink and DNA is in blue. The count of bicolored chromosomes is summarized in Figure 3. The dotted line shows the chromosome longest axis. Scale bar is 5 μm. Some chromosomes are shown twice, on the transverse and lateral view to better show the chromatid axis.

### Supplementary Text

#### Population genomics and evolution of reproduction in *Mesorhabditis*

##### 1 Population genetic structure under the *Mesorhabditis* life cycle

###### 1.1.1 General presentation and definition of parameters

The aim of the model is to predict population genomic patterns expected under the *Mesorhabditis* life cycle. We consider a single neutral locus with an infinite allele model (IAM) of mutation. We consider a subdivided population with *K* demes, each of size *N*. The total population is thus *N_T_* = *KN*. The life cycle is as follows (Figure 1 redrawn from the main text.). The generations are discrete and non-overlapping. Migration occurs before reproduction according to the island model at rate *m*. This simple population structure allows modelling the breeding structure of *Mesorhabditis* with high level of consanguineous mating: the lower the deme size and the migration rates, the higher the level of consanguineous mating.

**Figure 1:**
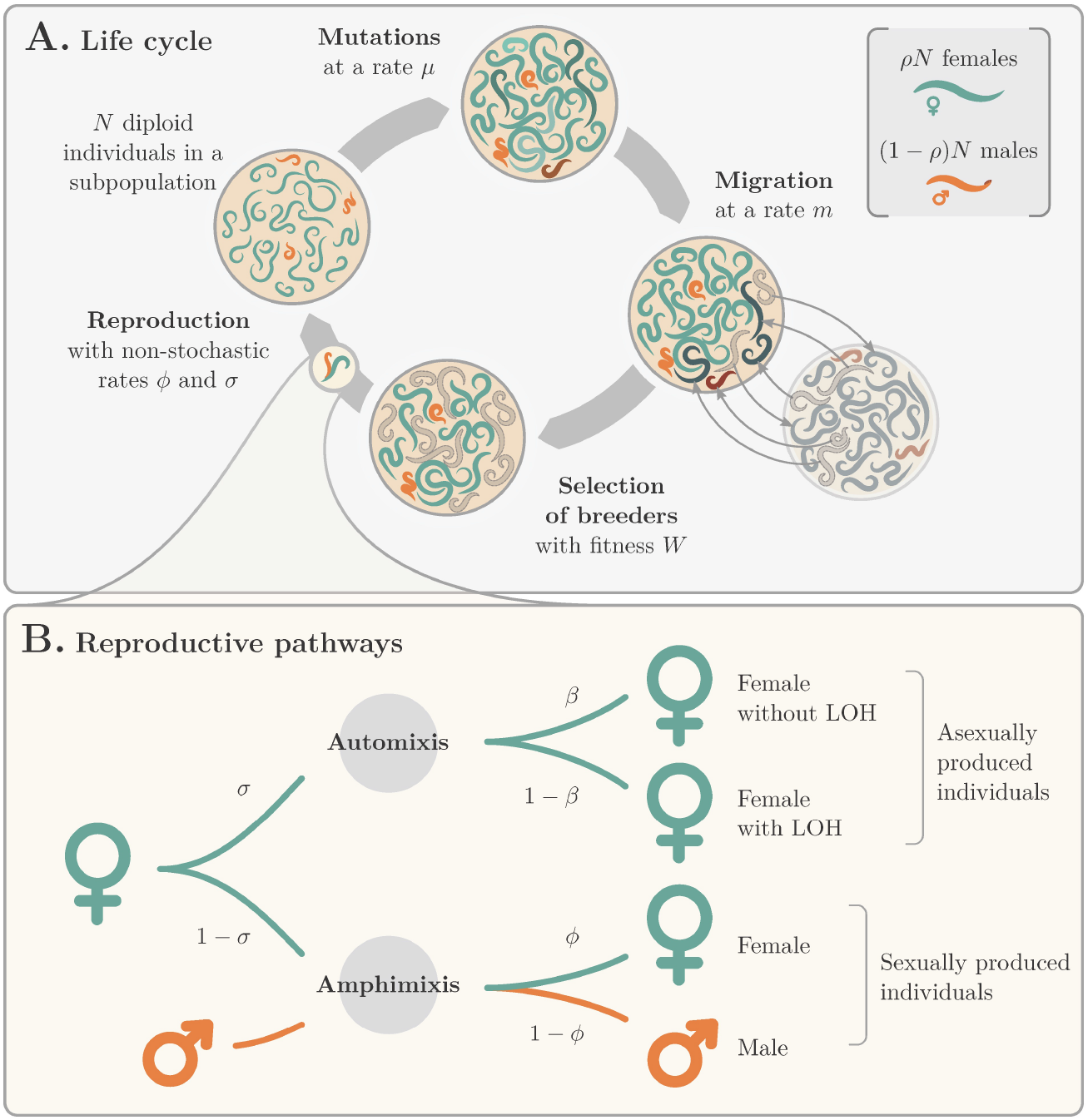
*Mesorhabditis* life cycle and parameters of the model.

Automictic reproduction that leads to gynogenetic females occurs in proportion *σ* and sexual reproduction 1 – *σ*. Under automixis, the rate of loss of heterozygosity (LOH) is *β* and depends on the underlying mechanism. With standard central fusion *β* = 1/2 after one crossover and random assortment of chromatids, and *β* = 0 if there is no recombination or in the proposed model of directed chromatid assortment (DCA, hereafter). Under sexual reproduction, the proportion of females and males is *ϕ* and 1 – *ϕ*, respectively. So far, no sexually-produced female has been observed (corresponding to *ϕ*=0). However, it is possible that they are produced at low rate in natural populations. If we set *σ* = 1 and *β* = 0, this is equivalent to a fully clonal model. If we set *σ* = 0 and *ϕ* = 1/2, this is equivalent to a fully sexual model.

To obtain measures of genetic diversity and population structure (F-statistics) we derive recursions on a set of probabilities of identity by descent (IBD), *Q_i_*, which leads to heterozygozity and F-statistics measure of the form [5]:

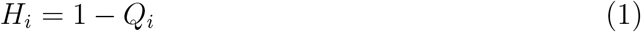

and

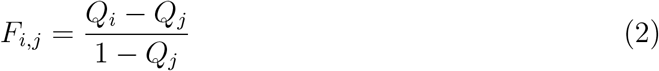

To fully describe the model, we need to follow eight probabilities of IBD, noted 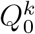 when the two genes are sampled in the same individual, 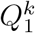 when they are sampled in two individuals of the same population, and 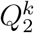, when they are sampled in two individuals from different populations. The superscript *k* stands for the sex of sampled individuals (see Figure 2).

**Figure 2:**
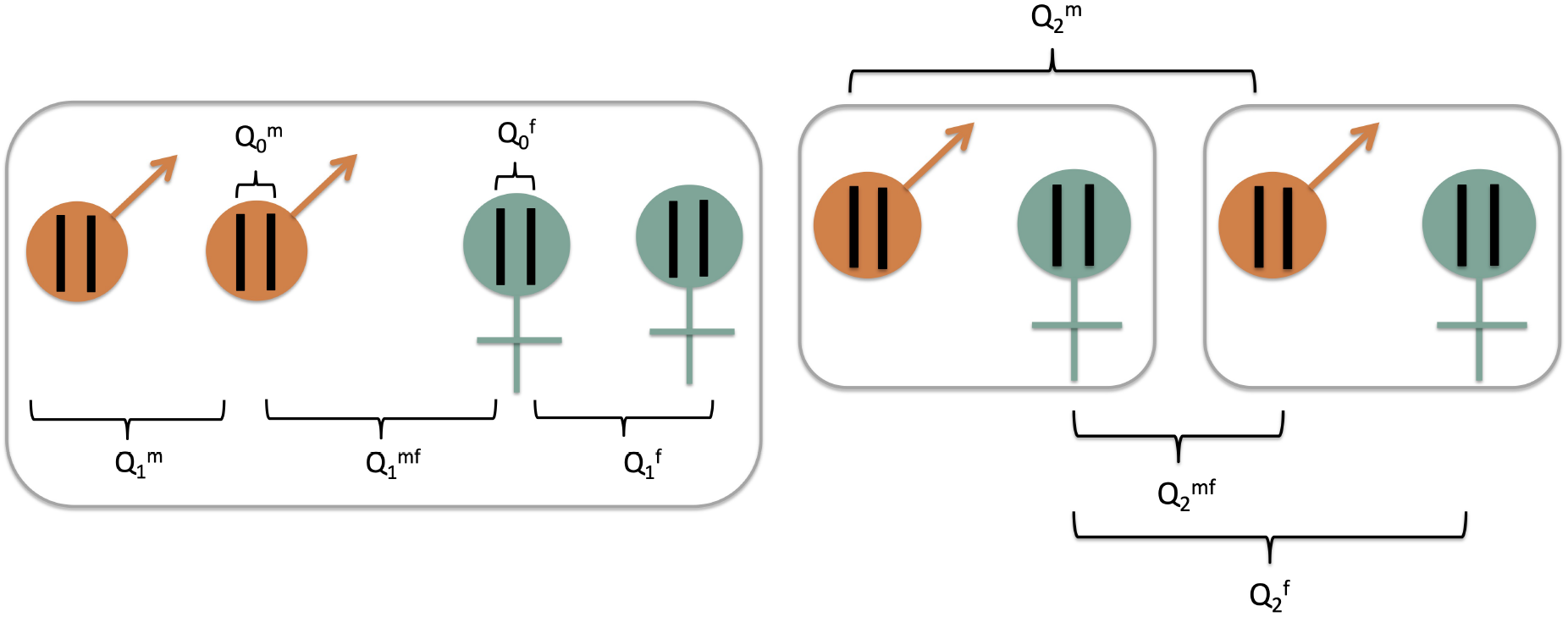
Definition of the eight probabilities of IBD. The grey boxes corresponds to demes.

From the parameters describing reproductive pathways (Figure 1) we obtain the proportion of males, sexual and gynogenetic females as:

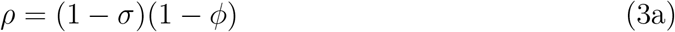

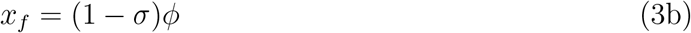

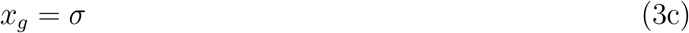

from which we obtain the number of males and females in the populations and the proportions of gynogenetic females among females

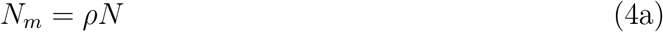

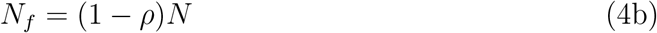

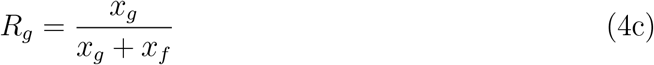

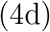

We also need to introduce the following compound parameters:

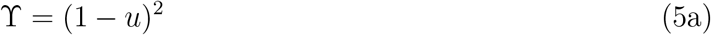

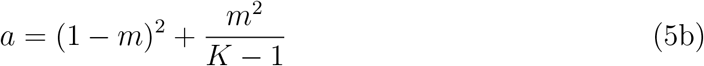

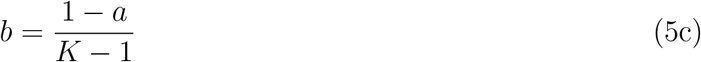

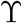 corresponds to the probability that the two sampled genes have not mutated in one generation, *a*, respectively *b*, corresponds to the probabilities that two individuals sampled in a same population, respectively in two different populations, were in the same population before migration.

###### 1.1.2 Recursions

We can now write the recursions of the *Q_i_*(*t* + 1) as a function of the *Q_i_*(*t*). At equilibrium *Q_i_*(*t* + 1) = *Q_i_*(*t*) so we remove the time subscript and directly give the equations at equilibrium:

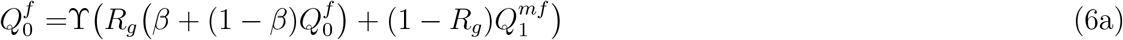

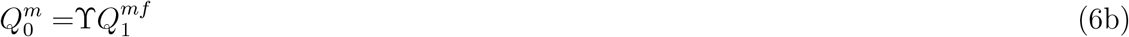

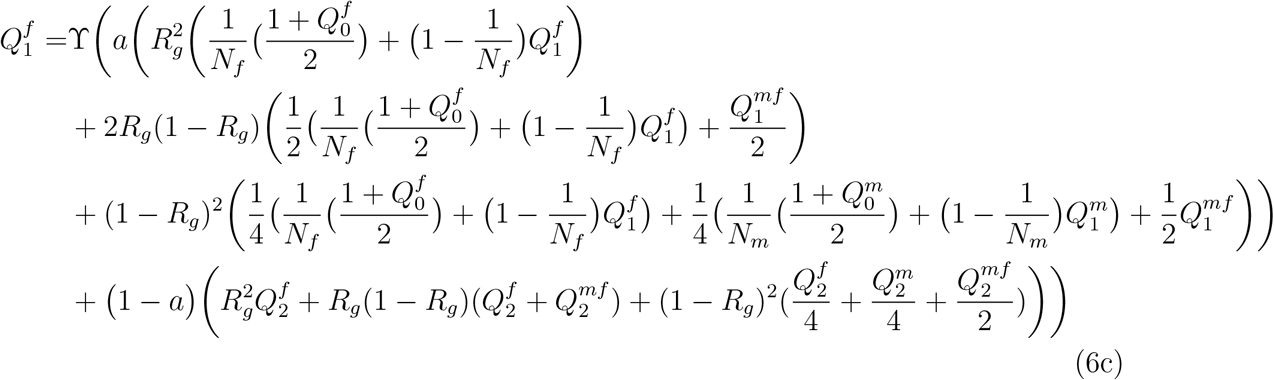

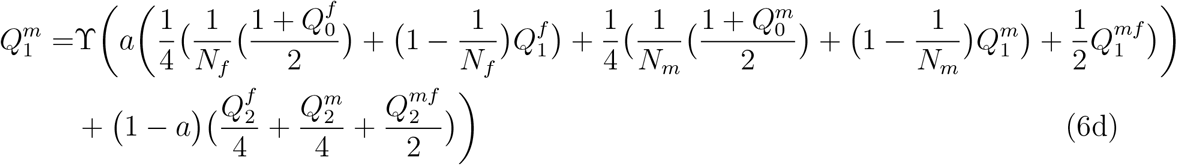

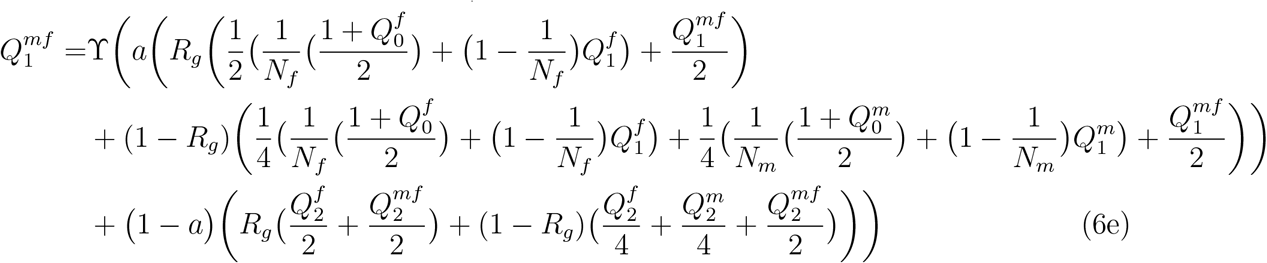

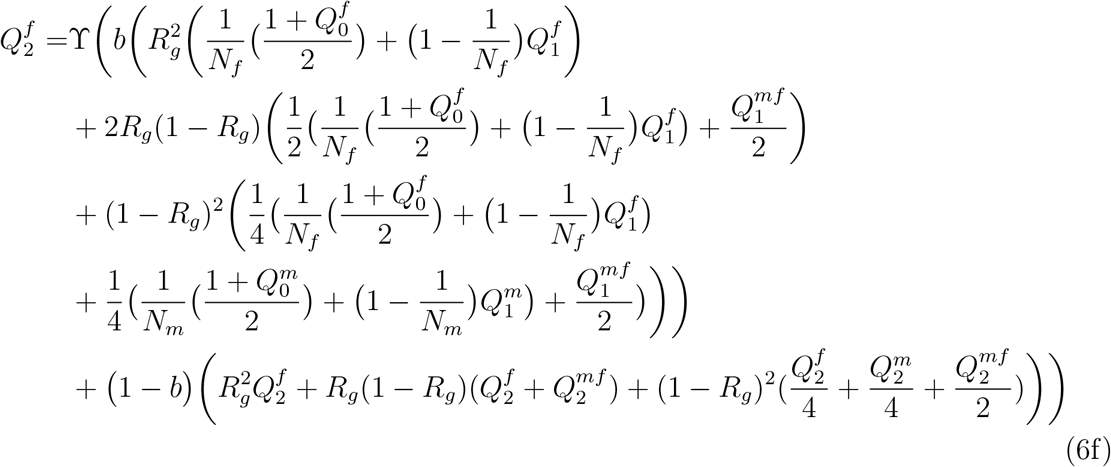

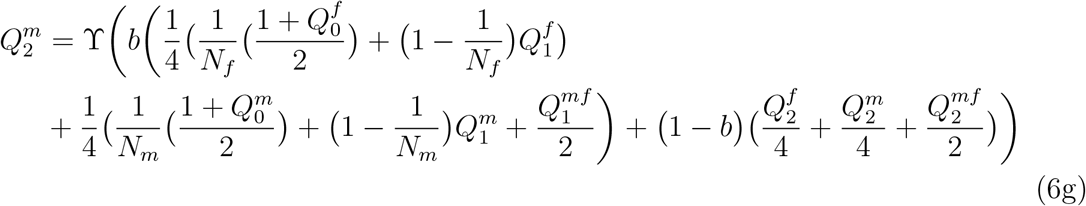

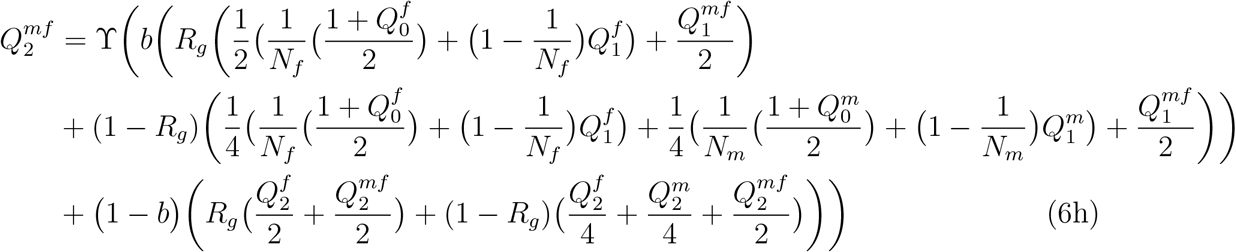

As an example, the rationale of the derivation is given for 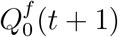. The two copies sampled in a female are IBD first if none has mutated 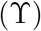. Then we must consider that this female is gynogenetic (*R_g_*) or sexual (1 – *R_g_*). If it is gynogenetic, if heterozygozity has been lost (*β*) the two copies are IBD with probability one, otherwise (1 – *β*) the probability of IBD is the same as for the mother, so 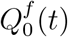. If the female come from sexual reproduction, the two copies are IBD with the same probability as of a random male/female pair at the previous generation, 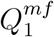. The terms in 1/*N_m_* and 1/*Nf* that appear in equations for the 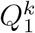 and 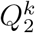 correspond to the probability that two different individuals have the same father or mother, respectively.

This system of recursions can be written in the matrix form:

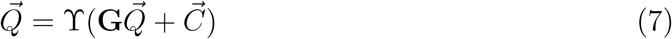

where 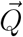 is the vector of probabilities of IBD,**G** is a matrix and 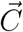 a vector, both depending of the parameters of the model. The solution can be written on the form:

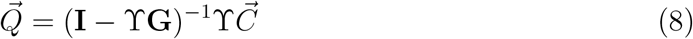

where **I** is the identity matrix.

The *F_IS_* statistics measured on genomic data corresponds to *F*_0,2_ in our model as we compare the IBD of two gene copies sampled either within an individual or at random over the whole population. It can be defined either for males, females, or for the whole population by weighting as a function of the sex-ratio:

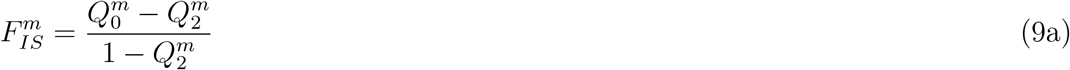

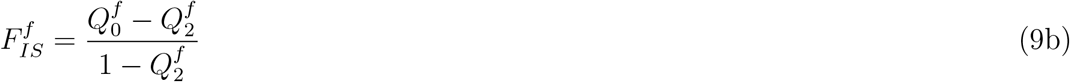

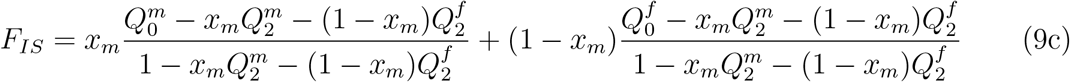

###### 1.1.3 Simulations

In addition to analytical derivations, an individual-based, multi-locus model was implemented using the SLiM software [3], where each individual’s genome is explicitly defined. It allowed to simulate genomic patterns along a chromosome and to introduce deleterious mutations in the model. Each individual is represented by a unique pair of autosome. This chromosome consist of *L* = 10^5^ loci, with a rate of recombination per locus *r* equal to 1/*L* = 10^-5^, so that on average one recombination event is observed per gametogenesis. Since *M. belari* has holocentric chromosomes (there is no specific centromere), the location of the chiasma is randomly drawn according to a uniform distribution along the chromosome. Two types of mutations are introduced, either neutral (no effect on fitness) or deleterious ones, with selection coefficient s, set to 0.01, and dominance coefficient *h*, set to 0.25, and acting multiplicatively across the genome. The life cycle was the same as described above. Various population sizes and migration rates were explored to modulate the level of inbreeding and we specifically studied the effect of the rate of production of sexual females, *ϕ* and the rate of LOH, *β*.

By adjusting the parameters of the model, we also compared the M. *belari* life cycle with more standard reproductive modes: full sexuality (*σ* = 0 and *ϕ* = 1/2), full clonality (*σ* =1 and r = 0), and central fusion automixis (*σ* = 1, and *β* = 1/2).

From the simulations we computed *F_IS_* and the linkage-disequilibrium measure *r*^2^ on neutral mutations at the scale of the whole population:

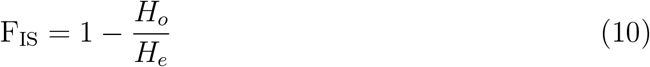

with *H_o_* the observed heterozygosity (percentage of heterozygous sites) and *H_e_* the expected heterozygosity:

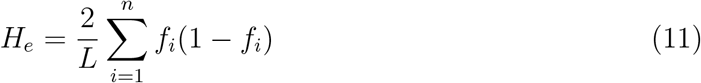

a sum over all the *n* mutations present in the population with their respective frequencies.

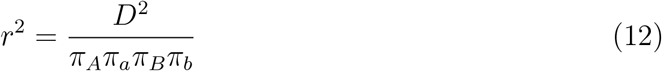

where *D* = *π_AB_* – *π_A_π_B_* with *π_A_* and *π_a_* the allelic frequency at a first locus *A* and *π_B_* and *π_b_* at a second locus *B*. Only mutations in frequency higher than 5% in the population were used. *r*^2^ was calculated for pairs of loci as the function of their distance on the simulated chromosome.

###### 1.2.1 F_IS_

The general solution of equation (8) can be obtain with the help of *Mathematica* [6] but it is formidable, so useless for direct biological interpretation. We thus performed numerical explorations in the general case. We also obtained approximations under two limit conditions: *ϕ* = 0 (no sexually-produced females) and *β* = 0 (no LOH). Under these two conditions we also used the standard diffusion limit. First we used the following scaling parameters: *θ* = 4*N_T_u*, Φ = 2*N_T_ϕ, B* = 2*N_T_β*. Then, we assumed an infinite number of local populations so *K, N_T_* → ∞ but that the scaled parameters terms tends towards constant: *θ*, Φ, *B* → *cte*. We also noted *M* = 4*Nm*, where migration is scaled with the local, *N*, which can be small.

Assuming no sexually-produced females we obtained:

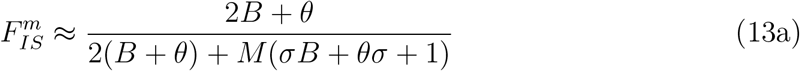

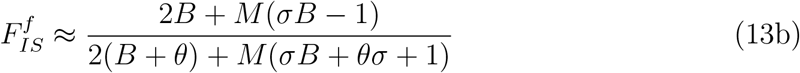

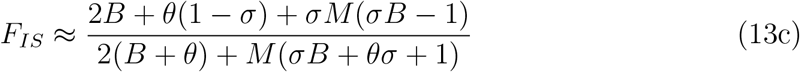

Assuming no LOH we obtained:

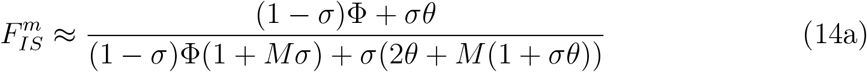

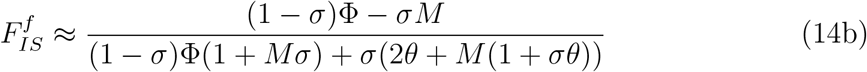

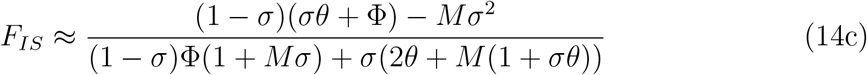

**Figure 3:**
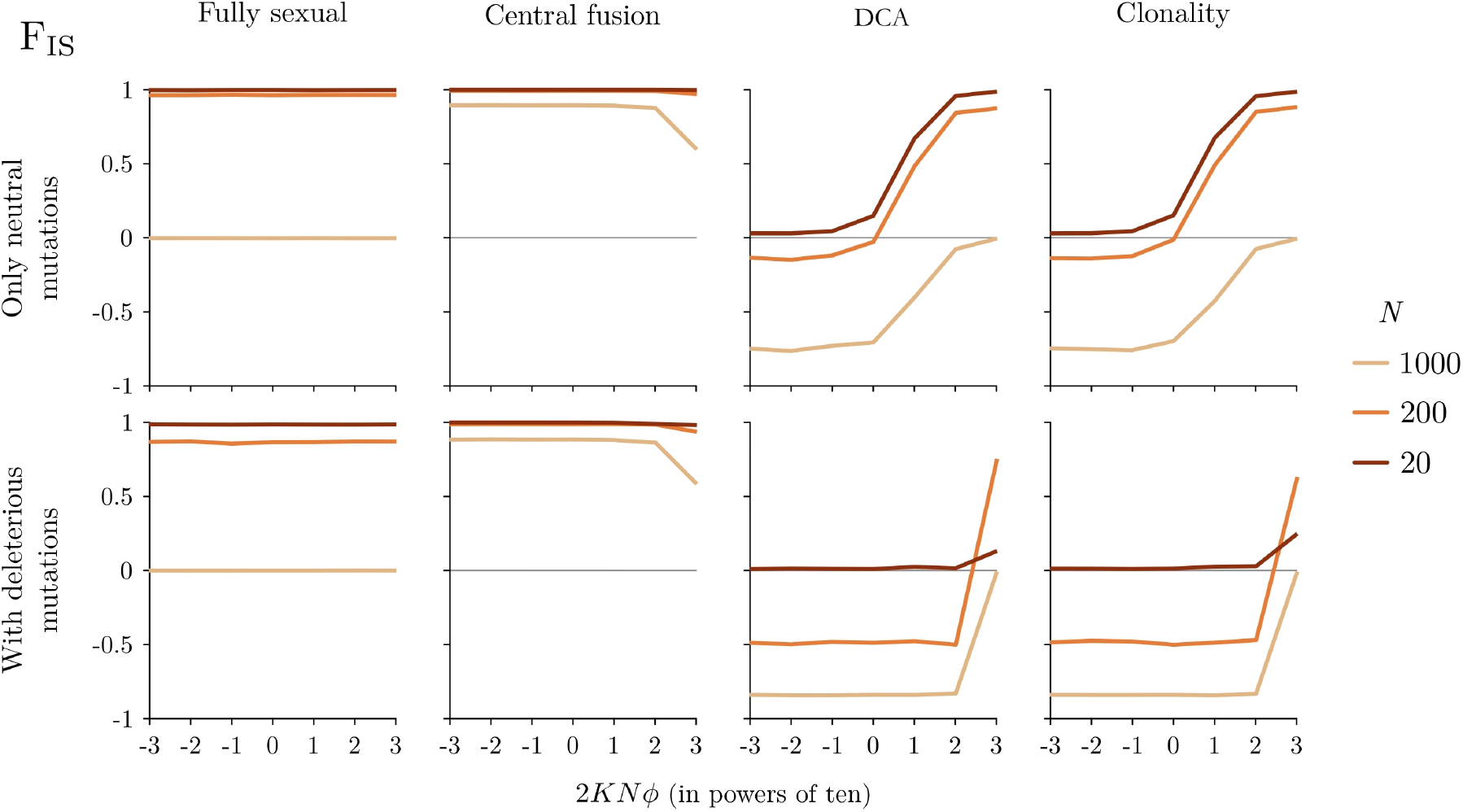
*F_IS_* for the different reproductive modes as a function of the rate of sexually produced females (2*KNϕ*) for different levels of population structure (*KN =* 1000, so *N* = 1000 corresponds to a single populations and *N* = 20 to a highly structured one).

We give the three expressions for completeness but we can concentrate on the expression for the average *F_IS_* In natural populations, there is a strong family structure with most matings occurring between kins. In our model, it corresponds to low migration between demes, so *M* close to 0. Then, equations (13c) and (14c) become:

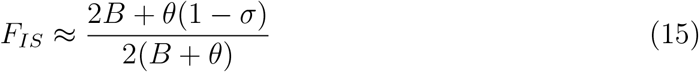

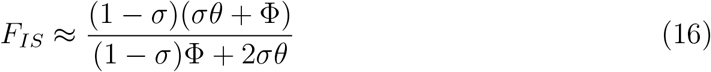

which both reduce to *F_IS_* = (1 – *σ*)/2 when *B* = 0 or Φ = 0. As *σ* ≈ 0.9 in natural populations it corresponds to *F_IS_* ≈ 0.05, so close to the observed value (*F_IS_* ≈ 0.19, see main text). In an unstructured population (*M* → ∞, and see *N* = 1000 on figure 3), *F_IS_* tends to −1 with no LOH (*B* = 0) (see also [1]). Here, this is compensated by the strong family structure leading to 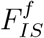 close to 0 as illustrated on figure 3 (see also [2]). It is worth noting that just a little bit of sex or LOH (higher than the mutation rate: *ϕ,B > θ*) rapidly leads to 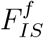 close to 1. This is confirmed by simulations as presented (3 and Figure 4 in the main text). This result is thus very sensitive to the occurrence of sexually produced females and to low level of LOH. For comparison,

However, simulations showed that because of deleterious mutations, the range of parameters leading to 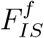 close to 0 can be much wider, even when some females are sexually produced and when the DCA is not complete (Figure 4). This is explained by the selection against highly homozygotes individuals. This is an important result as a non-zero proportion of sexually produced females is required to explain the low level of linkage disequilibrium observed genome wide as presented below.

**Figure 4:**
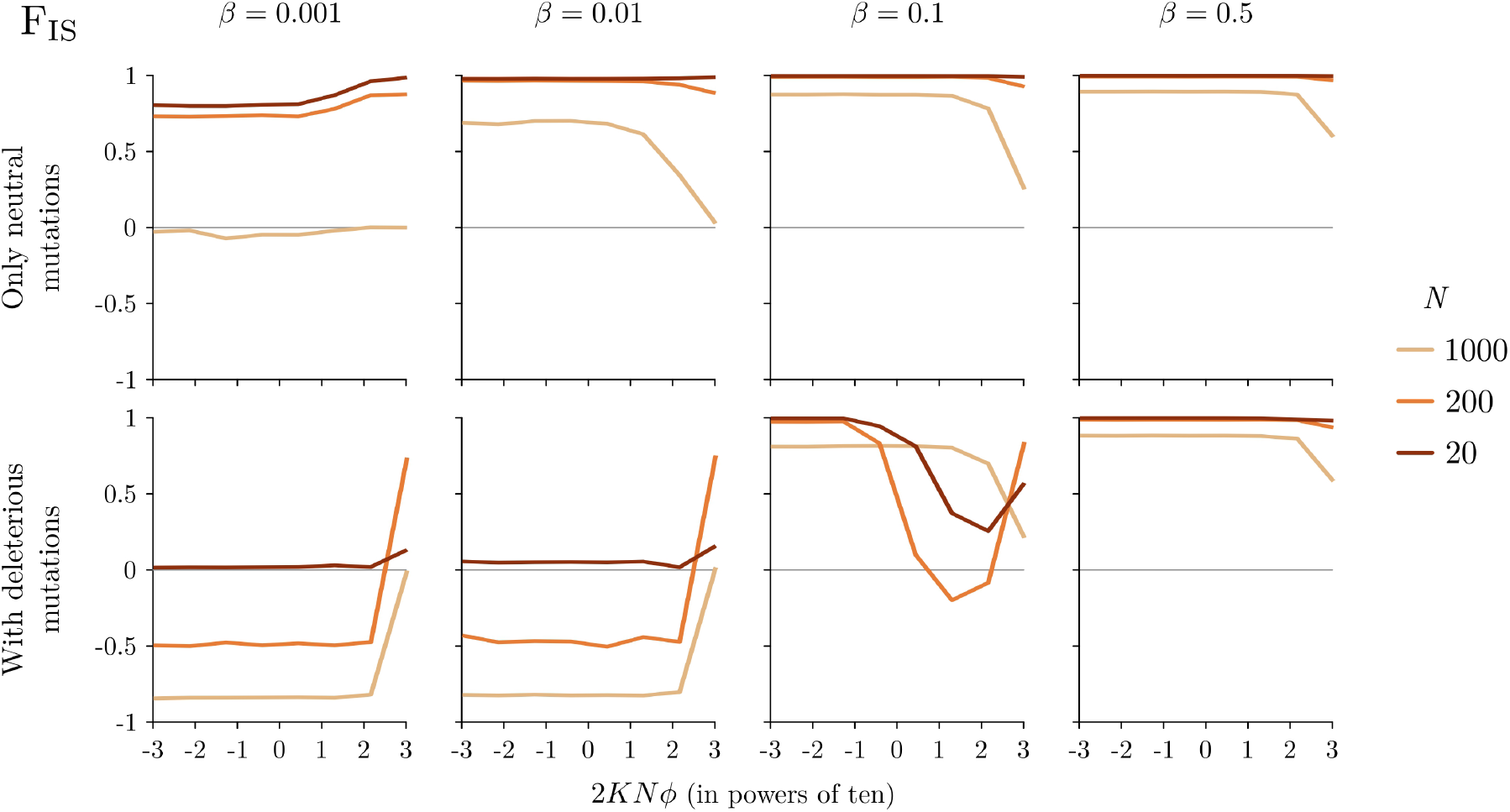
*F_IS_* for automixis with different level of DCA (β = 0 corresponds to full assortment and *β* = 1/2 to random pairing).

For comparison, figures 3 and 4 also shows the results for full sexuality, full clonality and standard central fusion automixis (without DCA).

###### 1.2.2 Linkage disequilibrium

Analytical derivations for linkage disequilibrium would require recursion equations for 40 IBD probabilities. We thus only relied on simulations.

**Figure 5:**
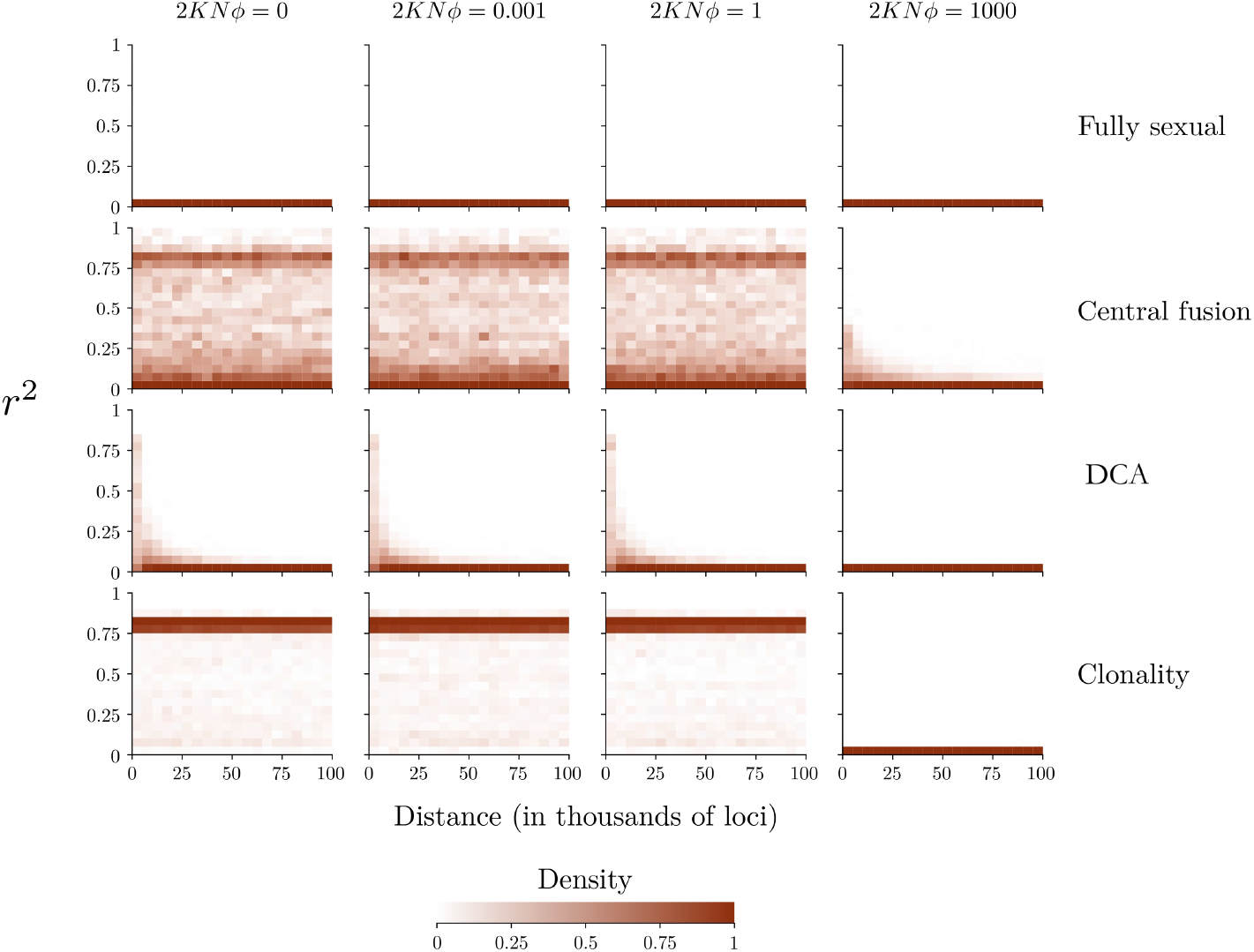
Pattern of linkage disequilibrium as a function of physical distance (in number of loci) for different rates of sexually produced females (2*NKϕ* = 0.001,1,100) and for different reproductive modes.

To understand the specific effect of the DCA we first considered a single unstructured population. As expected, LD is low and rapidly decreases with physical distance under sexual reproduction (Figure 5). In contrast, LD is high and independent of physical distance along chromosome under both full clonality because recombination does not occur, and under standard central fusion because recombination is not efficient as individuals are mostly homozygotes. Under complete DCA and without sexually produced females, LD pattern is intermediate: much lower than under clonality or standard central fusion but higher than under full sexuality, and decreasing with physical distance (Figure 5). The reason is that, although new haplotypes are generated by recombination within individuals, they are never associated together through mating, so recombination is not as efficient as under full sexuality.

When we also consider the effect of population structure, LD is globally higher and the difference among reproductive modes are less clear under pure neutrality. However, when deleterious mutations are added the differences become stronger. In particular a very low rate of sexually produced females is sufficient to make DCA similar to full sexuality whereas much higher rates are necessary to erase the signature of clonality or standard central fusion (Figure 6).

**Figure 6:**
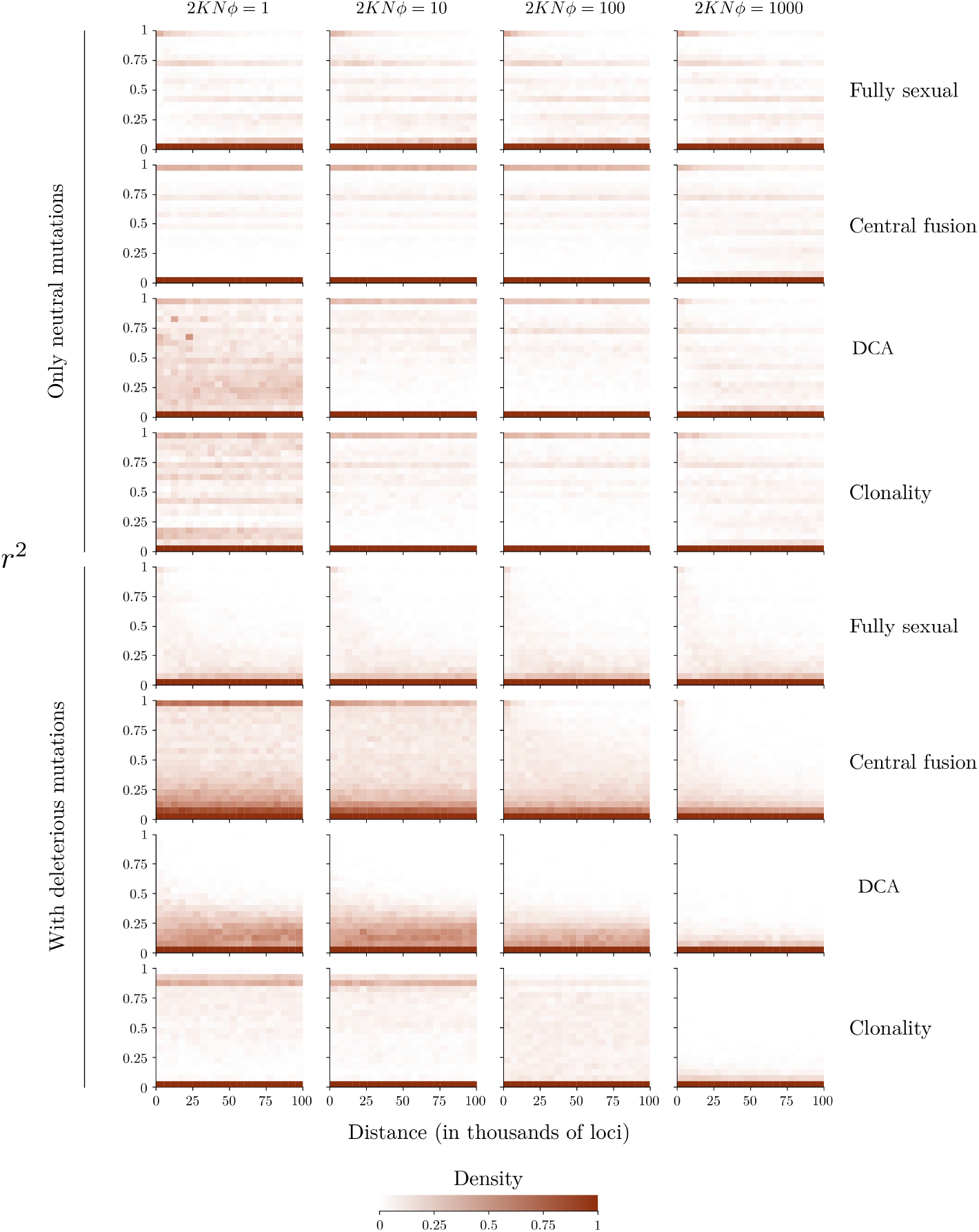
Pattern of linkage disequilibrium as a function of physical distance (in number of loci) for different rates of sexually produced females (2*NKϕ* = 0.001,1,100) and for different reproductive modes.

###### 1.2.3 Conclusion

Overall, the observed genomic pattern is well predicted by the strong family structure of *M. belari* populations directed chromatif assortment, sexually produced females at low rate (which can be unobserved in natural conditions), and the occurrence of deleterious mutations genome wide.

##### 2 Evolution of the reproductive system

In the second model, we study the evolution of the atypical reproductive system of *M. belari* from a standard sexual population. Compared to the previous model, the parameters σ (proportion of asexually produced offspring) and *β* (segregation bias) are no longer fixed but under the control of two independent evolving loci. Among sexually produced individuals the proportion of females is set to *ϕ* =1/2. Additional simulations where the sex-ratio can also evolve do not change the results and a sex ratio biased towards males evolved in a second step (not shown).

We started with a burn-in period where the population evolved under sexual reproduction. Then mutations are introduced at one or at the two loci controlling the reproductive mode. Each evolving locus is unlinked with any other locus. Transmission from parents to children of an evolving locus follows the same rules as transmission of loci on the focal chromosome, taking into account the potential loss of heterozygosity through chromosome segregation during gynogenesis. It can take two different values, a resident value, *g_r_*, or a mutant value, *g_m_* = *g_r_* ± δ. We used *δ* = 0.1 to avoid too long simulations. We assumed additivity between the two alleles so the genotypic value is the sum of the two alleles. At the beginning of simulations, all individuals are homozygous for the resident allele and a mutant allele is randomly introduced in one individual of the population. If this mutant becomes fixed in the population, this mutant allele becomes the resident allele. If the mutant disappears, another mutant is directly reintroduced into the population at random.

###### 2.1 Genotype to phenotype map

The value of σ is constrained between 0 and 1 and the value of *β* between 0 and 1/2. The gynogenesis rate starts at 0 and can theoretically go up to 1, the segregation bias and the segregation bias start at 0.5 and can reach 0 or 1. So, to map allelic values from] – ∞, +∞[ to [0,1] we used the following functions:

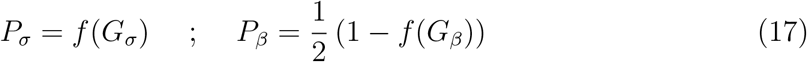

with

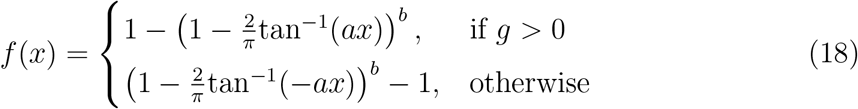

The parameters *a* and *b* define whether the function tends more or less quickly to 1 as x increases. We used *a* = 0.1 and *b* =15 but equilibrium results are not sensitive to chosen values. The gynogenesis rate and the segregation bias are controlled by the maternal genotype.

###### 2.2 Inbreeding depression

The supposedly strong barrier to the evolution of asexuality (assuming that automixis mechanisms can emerge) is inbreeding depression (ID) due to LOH that leads to the exposure of recessive deleterious alleles. It is calculated as 1 – *w*_1_/*w*_0_ with *w*_0_ the fitness of sexually produced individuals and w_1_ the fitness of asexually produced individuals.

A strong ID should hold back the evolution of automixis if the chromosome segregation bias is not strong, preventing LOH. Since we only modelled a single chromosome and central fusion leads to a LOH in only half of offsrpings, the maximum ID is 50%, which is the minimum value preventing the evolution of asexuality. Asexuality should always evolved under such a conditions. To allow a broader range of ID values we assumed an additional cost of LOH and corrected *w*_1_ to 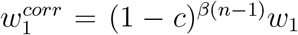, where *n* is the number of chromosomes (fixed to 10) and *c* is the mean fitness decline caused by LOH per chromosome.

###### 2.3 Extinction-recolonisation cycle of subpopulations

In this model, a higher migration rate is set, allowing the evolving loci to be transmitted not too slowly from one population to another. It should also be noted that here the parameters controlling reproduction are no longer fixed rates. It is thus possible that populations may stochastically run out of males, especially if gynogenesis evolves and the percentage of males decreases. If a population has no more males, it becomes extinct. An extinct population is recolonised in the generation following its extinction by a male and at least one female randomly selected from the other population. This pair reproduces and together they form the next generation of the subpopulation.

## Notes

### Competing Interest Statement

The authors have declared no competing interest.

